# The Cultural Brain Hypothesis: How culture drives brain expansion, underlies sociality, and alters life history

**DOI:** 10.1101/209007

**Authors:** Michael Muthukrishna, Michael Doebeli, Maciej Chudek, Joseph Henrich

## Abstract

In the last few million years, the hominin brain more than tripled in size. Comparisons across evolutionary lineages suggest that this expansion may be part of a broader trend toward larger, more complex brains in many taxa. Efforts to understand the evolutionary forces driving brain expansion have focused on climatic, ecological, and social factors. Here, building on existing research on learning, we analytically and computationally model the predictions of two closely related hypotheses: The Cultural Brain Hypothesis and the Cumulative Cultural Brain Hypothesis. The Cultural Brain Hypothesis posits that brains have been selected for their ability to store and manage information, acquired through asocial or social learning. The model of the Cultural Brain Hypothesis reveals relationships between brain size, group size, innovation, social learning, mating structures, and the length of the juvenile period that are supported by the existing empirical literature. From this model, we derive a set of predictions—the Cumulative Cultural Brain Hypothesis—for the conditions that favor an autocatalytic take-off characteristic of human evolution. This narrow evolutionary pathway, created by cumulative cultural evolution, may help explain the rapid expansion of human brains and other aspects of our species’ life history and psychology.

---

In the last few million years, the cranial capacity of the human lineage dramatically increased, more than tripling in size (Bailey & Geary, 2009; Schoenemann, 2006; Striedter, 2005). This rapid expansion may be part of a gradual and longer-term trend toward larger, more complex brains in many taxa (Balanoff, Bever, Rowe, & Norell, 2013; Dunbar & Shultz, 2007; Roth & Dicke, 2005; Shultz & Dunbar, 2010a; Striedter, 2005). These patterns of increasing brain size are puzzling since brain tissue is energetically expensive (Aiello & Wheeler, 1995; Elia, 1999; Foley, Lee, Widdowson, Knight, & Jonxis, 1991; Isler & Van Schaik, 2006; Kotrschal et al., 2013; Lieberman, 2011). Efforts to understand the evolutionary forces driving brain expansion have focused on climatic, ecological, and social factors (Bailey & Geary, 2009; Dunbar, 2003; Schoenemann, 2006; Striedter, 2005; van Schaik & Burkart, 2011). Here we provide an integrated model that attempts to explain both the broader patterns across taxa and the human outlier. To do this, we develop an analytic model and agent-based simulation based on the Cultural Brain Hypothesis (CBH): the idea that brains have been selected for their ability to store and manage information via some combination of individual (asocial) or social learning (Henrich, 2016; Heyes, 2012; Muthukrishna, 2015; Muthukrishna & Henrich, 2016; Reader, Hager, & Laland, 2011; Whiten & Van Schaik, 2007). That is, we develop the idea that bigger brains have evolved for more learning and better learning. The information acquired through these various learning processes is locally adaptive, on average, and could be related to a wide range of behavioural domains, which could vary from species to species. The forms of learning we model could plausibly apply to problems such as finding resources, avoiding predators, locating water, processing food, making tools, and learning skills, as well as to more social strategies related to deception, coercion, manipulation, coordination or cooperation. Our theoretical results suggest that the same underlying selective process that led to widespread social learning (Hoppitt & Laland, 2013) may also explain the correlations observed across species in variables related to brain size, group size, social learning, innovation, and life history. Moreover, the parameters in the formal representation of our theory offer hypotheses for why brains have expanded more in some lineages than others (Dunbar & Shultz, 2007; van Schaik, Isler, & Burkart, 2012).

Building on the Cultural Brain Hypothesis, our theoretical model also makes a set of predictions that we call the Cumulative Cultural Brain Hypothesis (CCBH). These predictions are derived from the parameters within that CBH model that favor an autocatalytic take-off in brain size, adaptive knowledge, group size, learning, and life history characteristic of human evolution. The CCBH has precedents in other models describing the processes that led to human uniqueness (see Boyd & Richerson, 1996; Dean, Kendal, Schapiro, Thierry, & Laland, 2012; Dean, Vale, Laland, Flynn, & Kendal, 2014; Henrich, 2016; Henrich & McElreath, 2003; Herrmann, Call, Hernández-Lloreda, Hare, & Tomasello, 2007; Heyes, 2012; Lewis & Laland, 2012; Reader et al., 2011). Since the CCBH is not a separate model, but instead additional predictions derived from the CBH model, this approach both seats humans within the broad primate spectrum created by the selection pressures we specify, and also accounts for our peculiarities and unusual evolutionary trajectory. That is, the same mechanisms that lead to widespread social learning can also open up a novel evolutionary bridge to a highly cultural species under some specific and narrow conditions—the CCBH. When these conditions are met, social learning may cause a body of adaptive information to accumulate over generations. This accumulating body of information can lead to selection for brains better at social learning as well as storing and managing this adaptive knowledge. Larger brains, better at social learning, then further foster the accumulation of adaptive information. This creates an autocatalytic feedback loop that enlists sociality, social learning, and life history to drive up both brain size and adaptive knowledge in a culture-gene co-evolutionary duet—the uniquely human pathway. The juvenile period expands to provide more time for social learning. As biological limits on brain size are reached (e.g. due to difficulties in birthing larger brains, even in modern populations, see Lipschuetz et al., 2015), increases in the complexity and amount of adaptive knowledge can take place through other avenues, such as division of information (and ultimately, division of labor), mechanisms for increasing transmission fidelity, such as compulsory formal schooling, and further expansion of the “adolescent” period between fertility and reproduction, spent in additional education (i.e. delayed birth of first child). (Muthukrishna & Henrich, 2016). This process modifies human characteristics in a manner consistent with more effectively acquiring, storing, and managing cultural information.

The CBH and CCBH are related, and can be explored with the same model, but we keep them conceptually distinct for two reasons. First, the cumulative culture-gene co-evolutionary process produces cultural products, like sophisticated multi-part tools and food processing techniques, that no single individual could reinvent in their lifetime (despite having a big brain capable of potent individual learning; Henrich, 2016). The evolution of a second inheritance system—culture—is a qualitative shift in the evolutionary process that demands analyses and data above and beyond that required for the CBH. Second, it’s possible that either one of these hypotheses could hold without the other fitting the evidence—that is, it might be the CCBH explains the evolutionary trajectory of humans, but the CBH doesn’t explain the observed patterns in social learning, brain size, group and life history in primates (or other taxa); or, vice-versa.

Our approach is distinct, but related to the Social Brain Hypothesis (SBH; Dunbar, 1998), which argues that brains have primarily evolved for dealing with the complexities of social life in larger groups (e.g., keeping track of individuals, Machiavellian reasoning, and so on). Initial evidence supporting the SBH was an empirical relationship shown between social group size in primates and some measure of brain size (different measures of brain size are typically highly correlated; Dunbar, 2009). Though this relationship does not hold outside the primate order, broader versions of the SBH that encompass other aspects of social cognition have been informally proposed with corresponding evidence from comparative studies. For example, a relationship has been shown between brain size and regular association in mammalian orders (Shultz & Dunbar, 2010a; Shultz & Dunbar, 2007), mating structure in birds and mammals (Shultz & Dunbar, 2007), and social structure and behavioral repertoire in whales and dolphins (Fox, Muthukrishna, & Shultz, 2017). Efforts to formally explore these ideas isolate three distinct evolutionary mechanisms. First, McNally and collaborators have explored the Machiavellian arms race between cooperation and deception (McNally, Brown, & Jackson, 2012; McNally & Jackson, 2013). Second, Dávid-Barrett and Dunbar (2013) simulate a relationship between coordination costs and group size showing that more complex coordination (and therefore higher cognitive complexity) is required as group size increases. Finally, exploring a distinct third mechanism, Gavrilets and Vose (2006) simulate an evolutionary competition among males for females in which males can evolve larger brains with learning abilities that permit them to acquire more effective strategies.

In his seminal paper, Humphrey (1976) highlighted the importance of social learning, along with several other social factors. The theory presented here is therefore consistent with this and other early research that emphasized the learning aspects of the social brain (Humphrey, 1976; Jolly, 1966; Whiten & Byrne, 1988a; 1988b; for a more recent discussion, see Whiten & van Schaik, 2007). However, while many verbal descriptions of the SBH are general enough to encompass most aspects of the CBH, formal instantiations of the SBH each focus on quite distinct evolutionary mechanisms: (1) deception and cooperation, (2) coordination between group members, and (3) learning social strategies. To make progress, we argue that it’s crucial to distinguish the various evolutionary mechanisms that have often been clumped under the “social brain” rubric, and then test for the action of these various mechanisms (which need not be mutually exclusive).

The CBH and CCHB are a deliberate shift in focus from “social” to “learning”; a shift with precedence in other theories, most informally expressed (for example, see Pradhan, Tennie, & van Schaik, 2012; Reader et al., 2011; Reader & Laland, 2002; van Schaik & Burkart, 2011; van Schaik et al., 2012; Whiten & Van Schaik, 2007). There are, however, some clear departures from most previous approaches. First, crucial to this shift from social to learning is that group size evolves endogenously, rather than as a product of externalities (such as avoidance of predators). Second, learning is assumed to be more general than the skills and cognition required for social living. Individuals could learn skills and knowledge for social coordination, cooperation, and competition, such as social strategies to improve mating, as in Gavrilets and Vose (2006). But equally, these skills and knowledge may be related to other fitness relevant domains, such as ecological information about finding food or making tools. Indeed, the generality of adaptive knowledge is critical to the CCBH and the human take-off. In our approach, the potential for a runaway process to explain the human outlier arises neither from a Machiavellian arms race (McNally et al., 2012; McNally & Jackson, 2013) nor from sexual selection (Gavrilets & Vose, 2006), but instead from the rise of cumulative cultural evolution as a second system of inheritance. Ecological factors are considered in the CBH in terms of survival returns on adaptive knowledge (e.g. easier acquisition of more calories or easier avoidance of predators, where easier means requiring less knowledge).

To further develop the CBH and CCBH, our models explore the interaction and coevolution of (1) learned adaptive knowledge and (2) genetic influences on brain size (storage/organizational capacity), asocial learning, social learning, and an extended juvenile period with the potential for payoff-biased oblique social learning. We explicitly model population growth and carrying capacity alongside genes and culture in order to theorize potential relationships between group size and other parameters, like brain size and adaptive knowledge, and also to examine the effects of sociality on the co-evolutionary process through two different parameters. We assume carrying capacity is increased by the possession of adaptive knowledge (e.g., more calories, higher quality foods, better predator avoidance). Our model incorporates ecological factors and phylogenetic constraints by considering different relationships between birth/death rates and both brain size and adaptive knowledge. This allows us to formalize (and in particular, simulate) these evolutionary processes for taxa facing diverse phylogenetic and ecological constraints.

## Models

We begin by laying out the key assumptions underlying both the analytical and simulation models. Then, using adaptive dynamics, we present our analytical model. From this model, we derive some key insights without the complexities of simulation. We then build on the analytic solutions to fully explore the mechanisms underlying these insights using an evolutionary simulation. This simulation also allows us to relax some of our assumptions, allowing oblique learning, learning biases, and life history to evolve and explicitly tracking group size.

We present the key insights and predictions of our model in three ways. First, we explain the conditions under which we expect relationships between our variables and how the size of these relationships is affected by our parameters. In doing so, we verbally describe the core logic underlying the theory. Second, we compare our predictions to existing data, plotting our simulation results side-by-side with this existing data. If our predictions were inconsistent with existing empirical correlations, this would pose a significant challenge to our theory. Finally, we derive the Cumulative Cultural Brain Hypothesis predictions, laying out the narrow evolutionary regime under which an autocatalytic interaction between cultural and genetic inheritance is most likely to generate a human-like take-off.

### Assumptions

Three key assumptions underlie our theory:

1. Larger and more complex brains are more costly than less complex brains because they require more calories, are harder to birth, take longer to develop, and have organizational challenges. Therefore, *ceteris paribus*, increasing brain size/complexity decreases an organism’s fitness. For simplicity, we assume that brain size, complexity, and organization(e.g., neuronal density) are captured by a single state variable, which we will refer to as“size”.
2. A larger brain correlates with an increased capacity and/or complexity that allows for the storage and management of more adaptive knowledge. Adaptive knowledge could potentially relate to locating food, avoiding predators, securing mates, processing resources(detoxification, increased calorie release), hunting game, identifying medicinal plants, making tools, and so on.
3. More adaptive knowledge increases an organism’s fitness either by increasing its number of offspring compared to conspecifics and/or by reducing its probability of dying before reproduction. Adaptive knowledge can be acquired asocially, through experience and causal reasoning, or socially, by learning from others.

The logic that follows from these key assumptions is first formalized using an analytic approach—an adaptive dynamics evolutionary model (Doebeli, Hauert, & Killingback, 2004). This model captures the logic and several of the key predictions of the CBH. We then simulate the logic to capture the co-evolutionary dynamics needed to generate the CCBH.

### Analytical Model

To explore the evolutionary adaptive dynamics of the CBH, we begin with individuals *i* represented by three continuous variables: brain size *b*_*i*_, adaptive knowledge *a*_*i*_, and reliance on social learning (over asocial learning; e.g. time spent), *s*_*i*_. We will initially ignore the evolution of oblique learning, learning biases, and population structure, and assume that individuals using social learning use oblique learning and learning biases to hone in on the target individual with the most adaptive knowledge. We will relax this assumption in our simulation and allow oblique learning, learning biases, life history, and population structure to endogenously evolve. Table 1 is a handy key for the variables in our analytic model.

**Table 1.**
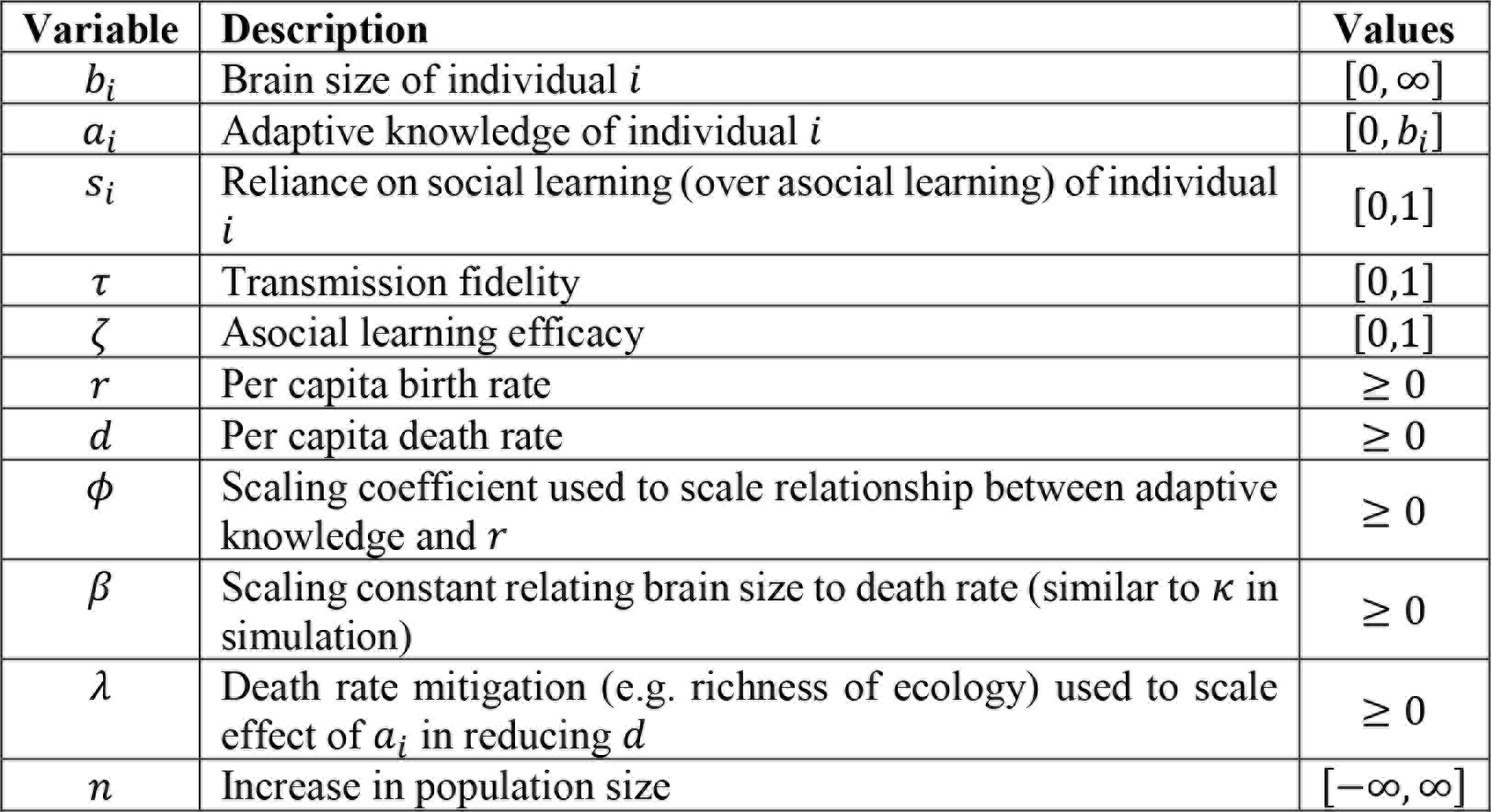
Variables for analytic model.

Individual *i* has two routes to acquire adaptive knowledge *a*_*i*_: (1) through asocial (individual) learning as a function of their own brain size *b_t_* and (2) through social learning as a function of the target model from whom they learn (*a*_*m*_). The proportion of time or propensity to use social over asocial learning is given by *s_i_*. Thus adaptive knowledge is given by:

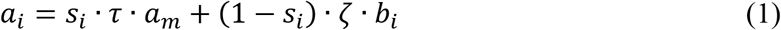

Where *τ* is transmission fidelity (how well an individual can learn from a model), *ζ* is asocial learning efficacy (how effectively an individual can use their brain to figure things out), and *a_m_* is the adaptive knowledge possessed by the individual in the parent generation from whom they are learning (e.g. model with maximal *a* or model with average *a*, etc).

The parameters τ and *ζ* in Equation 1 are abstractions of more complicated details covered in other work. By outsourcing the evolution of these features to other models, we can focus on the core of the CBH argument; i.e. how learning, brain size, knowledge, sociality, and life history are interconnected. Examples of this earlier works include, Lewis and Laland (2012) model of the relationship between transmission fidelity and the rate of trait loss, showing that sufficiently high transmission fidelity is necessary for cumulative culture, even more so than novel invention, incremental improvement, and recombination. Relatedly, building on work by Henrich (2004), Mesoudi (2011) models how increases in cumulative culture (driven by, for example, sociality) are more difficult for each generation to acquire. Thus, selection favors mechanisms to increase transmission fidelity. Muthukrishna and Henrich (2016) discuss the many mechanisms to increase transmission fidelity as adaptive knowledge accumulates. Mechanisms such as explicit teaching may not be required in a small-scale society, but in a large-scale society, not only is explicit teaching required, but also formal institutionalized schooling from a variety of teachers. Thus, *τ* could include individuals’ cognitive abilities (itself increased by culture; see Muthukrishna & Henrich, 2016), but also greater social tolerance, more interactions or opportunities for interaction, and some passive or active teaching by models, and so on (for more examples, see Dean et al., 2012; Muthukrishna & Henrich, 2016; Whiten & Erdal, 2012). Transmission fidelity could be broken down into constraints and endogenous state variables for genetic, cultural, and social factors, as well as interactions between these (e.g. genes for sociality), but for the purposes of expressing our argument, here we capture all this with *τ*.

Similarly, our model relies on the idea that “bigger” brains will be better at solving novel problems, and figuring stuff out (Deaner, Isler, Burkart, & van Schaik, 2007; Sol, Bacher, Reader, & Lefebvre, 2008). As Deaner et al. (2007) analyses reveal, at least in primates, the best predictor of cognitive ability is overall brain size. But, as with transmission fidelity, many factors will influence individuals’ ability to use their brains, such as constraints on time (for trial and error learning) or energy. These constraints are captured by *ζ*.

We take an evolutionary adaptive dynamics approach to find the evolutionary stable strategies (ESS) in our model. This approach involves assuming a monomorphic population and then looking at the “invasibility” of the population to a mutant (in variables of interest) with slightly different values. Appropriate to the dynamics we are interested in, this analytic method assumes mutations are small (i.e. we are not exploring competition between two vastly different groups).

#### Social Learning

To determine the average adaptive knowledge in a population that is monomorphic for resident genotype (*s, b*), we’ll initially assume that genotype is fixed over the course of learning. We’ll assume that the learning process leads to a distribution of adaptive knowledge values in the population and that individuals using social learning select a model using payoff-biased learning, choosing to learn from the model with the maximal possible value of adaptive knowledge *a_max_* (i.e., they learn from the rare individual who has attained the maximal value). In the simulation, we will relax this assumption and allow oblique learning (learning from non-genetic parents) and learning bias to evolve. Assuming individuals do learn from the best model when social learning, the mean adaptive knowledge in the population is given by:

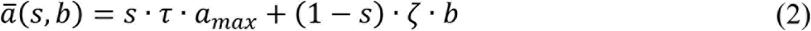

We further assume that the maximal adaptive knowledge is constrained by the brain size of the learner, such that *a*_*max*_ = *vb*, where *v* > 0 is some scaling parameter. As we shall see, the insights of the model are independent of the specific value of the *v* scaling. Thus, Equation 2 becomes:

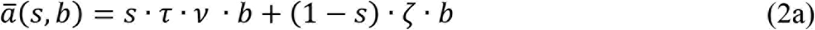

We can now easily understand the adaptive dynamics of the social learning trait (*s*) assuming more adaptive knowledge has a higher payoff. For a given brain size *(b*), we simply compare *τvb* and *ζb*: if *τvb* > *ζb*, then it pays to increase s as much as possible to maximize adaptive knowledge (i.e. *s* → 1); conversely, if *τvb* < *ζb*, then it pays to decrease *s* as much as possible to maximize adaptive knowledge (i.e. *s* → 0). This will be true as long as individuals have access to a range of models and are learning from the model with the greatest adaptive knowledge. Given these conditions, the key to reliance on social learning is the ability to learn with high fidelity and the key to reliance on asocial learning is the ability to efficiently use one’s brain to learn by oneself. Further, if there is some limitation on accessing the model with the maximal adaptive knowledge, such as ineffective payoff biased learning making it difficult to identify who has the most adaptive knowledge or too small or disconnected a population for at least one individual to consistently reach this maximal value every generation, then the evolution of social learning is also going to depend on the maximal adaptive knowledge learners have access to. We explore these dynamics in the simulation model.

#### Brain Size

To determine the adaptive dynamics of brain size, we need an ecological model for monomorphic populations (i.e. for populations that consist of a single resident type (*s,b*). To do this, we need to specify how the various traits affect the birth and death rates in the model. We use a logistic ecological model:

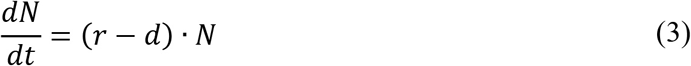

Here *N* is population density, *r* is the per capita birth rate of the resident and *d* is the per capita death rate of the resident. Next, we specify the per capita birth rate (*r*) and death rate (*d*). We assume the birth rate *r* decreases with population size (density dependence influencing carrying capacity), but that that decrease is slower with increased adaptive knowledge (e.g. allowing you to support more offspring or outcompete competitors in access to mating opportunities). The birth rate (*r*) is given by:

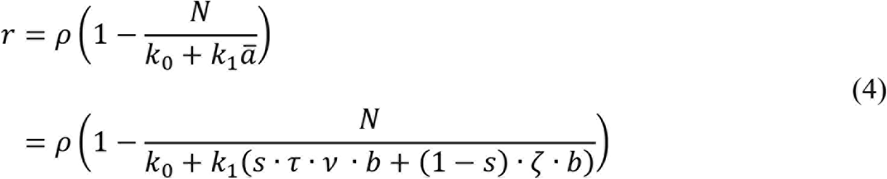

Where *ρ* is the maximal birth rate and that dependence leads to a linear decrease in the birth rate given by the second half of Equation 4. This linear decrease is assumed to be influenced by the mean adaptive knowledge (*a̅*), such that more adaptive knowledge leads to a larger denominator, slowing the decrease with density dependence (allowing for a higher effective carrying capacity). *k*_0_ and *k*_1_ are positive parameters, which we set to 1, without loss of generality, in the following analyses.

We assume that a larger brain is more costly than a smaller brain in terms of death rate (e.g. higher calorie requirements), but that more adaptive knowledge lowers the death rate (e.g. finding food or evading predators). The death rate (*d*) is given by:

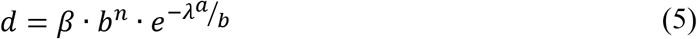

This function assumes that the cost of brains scales up in a polynomial fashion (e.g. *n* = 2), but that the reduction in the death rate through adaptive knowledge is an exponential decay, where adaptive knowledge is bounded by brain size (i.e. *a* ≤ *vb*). Here *β* scales the maximum brain size and *λ* scales the death rate reducing payoff to adaptive knowledge. The degree to which adaptive knowledge can offset brain size is a ratio of adaptive knowledge to brain size (adaptive knowledge is constrained by brain size regardless of learning mechanism and as brains grow, more knowledge is required to provide an equivalent offset) and *λ*. The *λ* parameter allows us to adjust the extent to which adaptive knowledge can offset the costs of brain size, where λ = 0 indicates no offset. The *λ* parameter can be interpreted as how much adaptive knowledge one requires to unlock the fitness-enhancing advantages. For example, in a calorie-rich environment where only a little skill or knowledge is required to access calories (e.g. simply remembering food locations), *λ* would be high. Conversely, in a calorie-poor environment where a lot of skills or knowledge are required to access fewer calories (e.g. food needs significant preparation before safe consumption), *λ* would be low. In the analytic model, the decrease to the death rate through adaptive knowledge becomes a constant since adaptive knowledge is a function of brain size (and parameters affecting learning efficiency), but although this will not affect the dynamics of the model, it will affect the final brain sizes. We fully explore this in the simulation.

Given a resident (*s, b*), the equilibrium population size of the resident is determined by the solution to Equation 3:

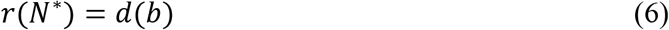

Since we know that *s* → 0 when *τν* < *ζ* and *s* → 1 when *τν* > *ζ*, we can consider these two cases, asocial learners and social learners, separately and then compare the outcomes of these two regimes.

#### Asocial learners (*s* = 0)

To determine the adaptive dynamics of brain size, consider a mutant (designated by subscript “m”) with brain *b*_*m*_. This mutant’s adaptive knowledge based on Equation 1 will be *a_m_* = *ζb_m_*, since *s* = 0. Using the same ecological assumptions as before for a mutant type *b_m_*, and assuming the mutant is rare and growing (initially) in a resident population that is at its ecological equilibrium *N*\* the per capita growth rate of the mutant, its invasion fitness, is:

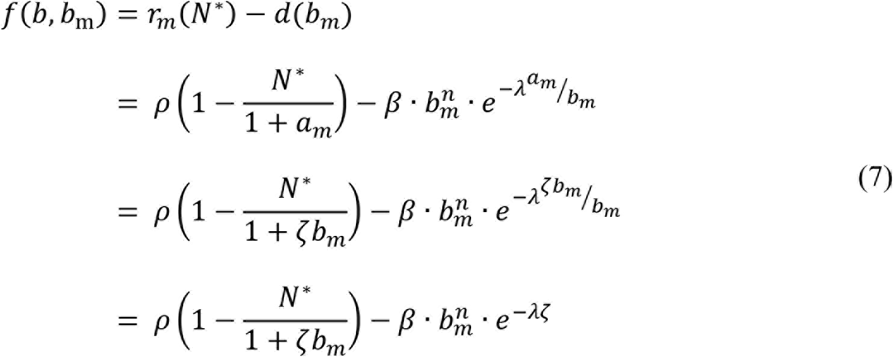

To examine the adaptive dynamics of brain size, we need to calculate the selection gradient by taking the derivative of the invasion fitness *f* with respect to the mutant trait *b_m_* and evaluate this derivative at the resident value *b*. To calculate if these equilibria are stable, we will calculate the second derivative. If the second derivative is negative, then the value is a convergent stable ESS. For those unfamiliar with this approach, it may be helpful to use a physical analog—distance, speed, and acceleration (or more accurately, displacement, velocity, and acceleration). The derivative of distance over time (metres) is speed (metres per second). The second derivative (derivative of speed) is acceleration (metres per second per second). The adaptive dynamics approach is the equivalent of looking at when an object is stationary (i.e. speed—derivative of distance—is 0) and confirming that these “equilibria” stationary points are convergent by confirming that objects decelerate around these points (i.e. acceleration—second derivative—is negative). If the second derivative were positive, objects would increase speed and move away from this stationary point, or in the present case, there would be positive selection for mutants away from this equilibrium. Let us calculate the selection gradient for brain size:

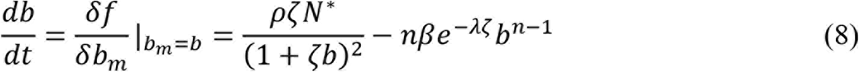

From Equation 8 we can see that if *n* > 1, *db/dt* < 0 for large *b* and *db/dt* > 0 for small *b*, which suggests that there is some intermediate ESS value for brain size *(b*\*). It is straightforward to check that the second derivative of the invasion fitness function (Equation 8) with respect to the mutant trait and evaluated at the resident trait is always negative and therefore the singular strategy *b*\* is a CSS (i.e., a convergent stable ESS). This equilibrium brain value (i.e. when *db/dt* = 0) is difficult to solve for a generic polymial *n*. To calculate a solution, we can select a reasonable polynomial (e.g. *n* = 2, which we use in the simulation) and solve for *db/dt* = 0. As long as brain size is positive, the relationship between brain size and the death rate will be superlinear and monotonous; our qualitative results should be robust to the specific polynomial used. Here is the equilibrium brain size for *n* = 2:

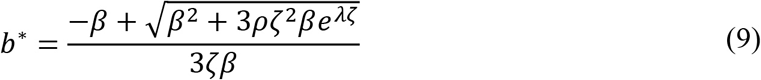

We need to compare the equilibrium brain size among asocial learners expressed in Equation 9 with the equilibrium brain size among social learners, so let’s now calculate the dynamics for social learners.

#### Social learners (*s* = 1)

To determine the adaptive dynamics of brain size, consider a mutant (designated by subscript “m”) with brain *b*_*m*_. This mutant’s adaptive knowledge based on Equation 1 will be *a*_*m*_ = *τvb*_*m*_, since *s* = 1. Using the same ecological assumptions as before for a mutant type *b_m_*, and assuming the mutant is rare and growing (initially) in a resident population that is at its ecological equilibrium *N*\*, the per capita growth rate of the mutant, its invasion fitness, is:

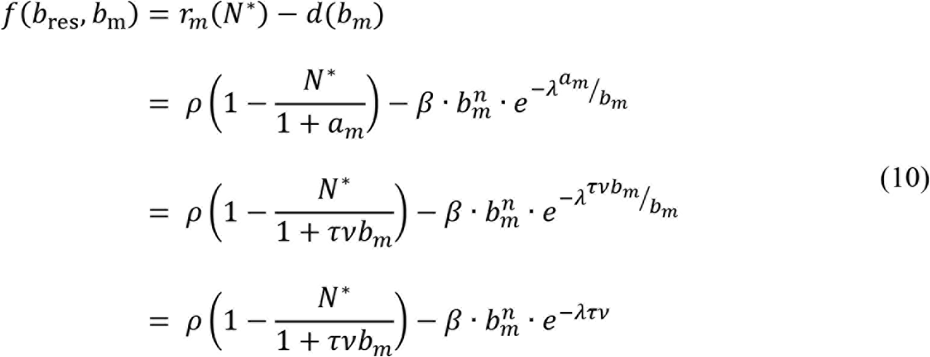

As before, to examine the adaptive dynamics of brain size, we need to calculate the selection gradient by taking the derivative of the invasion fitness *f* with respect to the mutant trait *b_m_* and evaluate this derivative at the resident value *b_res_*. To calculate if these equilibria are stable, we will calculate the second derivative. If the second derivative is negative, then the value is a convergent stable ESS. Let us calculate the selection gradient for the brain size of social learners:

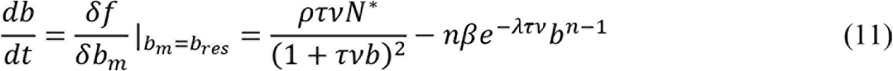

As with asocial learners, from Equation 11 we can see that if *n* > 1, *db/dt* < 0 for large *b* and *db/dt* > 0 for small *b*, which suggests that there is some intermediate ESS value for brain size (*b*\*). It is straightforward to check that the second derivative of the invasion fitness function (Equation 11) with respect to the mutant trait and evaluated at the resident trait is always negative and therefore the singular strategy *b*\* is a CSS (i.e., a convergent stable ESS). We can set *n* = 2 and calculate this equilibrium brain value (i.e. when *db/dt* = 0):

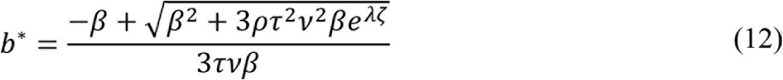

Equation 12 is functionally similar to Equation 9, but the equilibrium brain size for asocial and social learners will be different. Moreover, since to enter the realm of social learning, *τvb* > *ζb*, social learners, *ceteris paribus*, will have larger equilibrium brain sizes than asocial learners. However, transmission fidelity, asocial learning efficacy, and the payoff for adaptive knowledge (e.g. richness of the environment) are all going to affect the equilibrium brain size. We can derive a set of predictions from the insights gained from this model.

#### Predictions

The key predictions from the analytical model are that:

1. Increased reliance on social learning requires high transmission fidelity (relative to the ability to generate knowledge by oneself).
2. Extreme reliance on social learning also assumes access to a range of models with different amounts of adaptive knowledge (determined by population size and interconnectedness and assuming an ability to select and learn from models with more adaptive knowledge; see Henrich, 2004; Henrich et al., 2016; Muthukrishna & Henrich, 2016).
3. A greater return on adaptive knowledge (affected by *λ*; e.g. richness of environment) increases brain size (and may therefore explain different encephalization slopes across tax). Assuming an exponential return on adaptive knowledge, the environment will have a larger effect on social learners.

However, there are several assumptions and implications underlying these basic insights, such as:

1. Social learners face a bootstrapping problem of where the initial knowledge comes from.
2. The birth rate and the indirect relationships that affect actual population size will also affect brain size (and adaptive knowledge).
3. Species that do enter an extreme of social learning (such as humans) are on a treadmill, requiring higher transmission fidelity and more adaptive knowledge to sustain their large brains. A loss in either transmission fidelity or access to adaptive knowledge would drive the species towards a smaller brains.

Brain size and reliance on social over asocial learning will depend on factors that affect availability of adaptive knowledge, which are themselves affected by learning strategies and adaptive knowledge. In other words, there are a range of co-evolutionary dynamics that we have assumed or abstracted away in order to solve this model analytically, but which are crucial to capture and understand the full range of evolutionary dynamics. To understand the conditions under which social learning might emerge (and perhaps more interestingly, extreme reliance on social learning), we need to explore these co-evolutionary dynamics. We explore these full set of variables and explore these dynamics through an evolutionary simulation. An evolutionary simulation also allows us to properly account for population size, population structure, more sophisticated learning strategies, and life history. This model will bolster and expand on our analytic model and reveal the conditions where adaptive knowledge and brain size will increase.

### Simulation Model

To explore the culture-gene co-evolutionary dynamics, we constructed an evolutionary simulation that extends our analytic model. In our simulation, individuals are born, learn asocially or socially from their parent with some probability, potentially update by asocial learning or by socially learning from more successful members of their group during an extended juvenile period, migrate between demes, and die or survive based on their brain size and adaptive knowledge. Individuals who survive this process give birth to the next generation. We are mainly interested in the effects of natural selection and learning, so we use a haploid model and ignore non-selective forces such as sex, gene recombination, epistasis, and dominance. The lifecycle of the model, as well as all variables and parameters, are shown in Figure 1 below.

**Figure 1.**
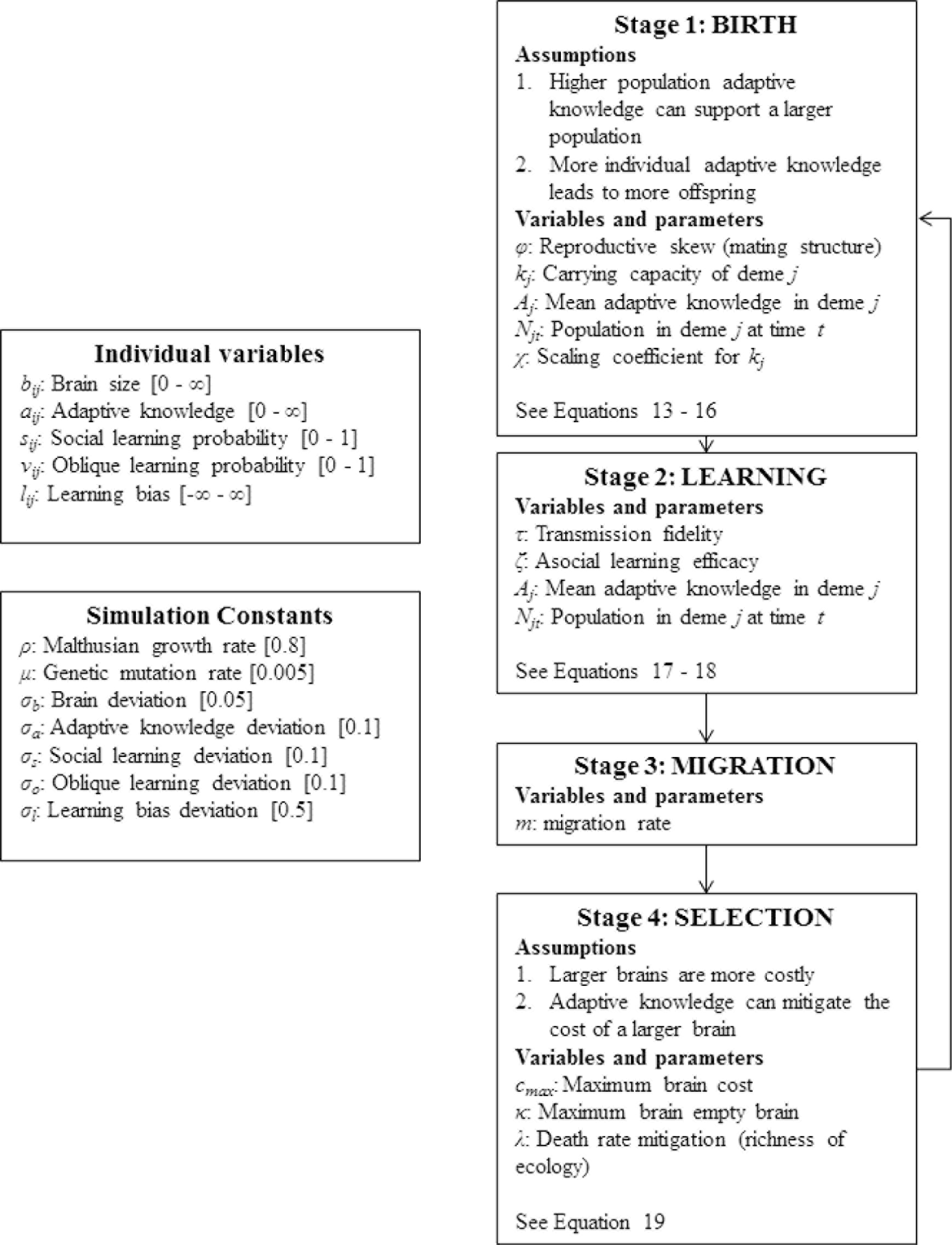
Lifecycle of simulation. On the left we define all individual evolving variables and constants. Parameters are defined within the relevant life stage.

This simulation was written in C++ by MM (code in Supplemental Materials). To reduce bugs, two computer science undergraduate research assistants wrote a suite of unit tests using Google’s C++ Testing Framework. These CS students also independently reviewed the code. The simulation begins with 50 demes, each with a population of 10 individuals. Throughout the simulation, the number of demes was fixed at 50. In early iterations of the model, we explored increasing the number of demes to 100 for some of the parameter space and found no significant impact on the results. Our starting population of 10 individuals is roughly equivalent to a real population of 40 individuals, assuming two sexes and one offspring per parent (4 × 10). As a reference, mean group size in modern primates ranges from 1 to 70 (Dunbar, 2009).

Each individual *i* in deme *j* has a brain of size *b_ij_* with a fitness cost that increases with increasing brain size. Adaptive knowledge is represented by *a_ij_*, where 0 ≤ *a_ij_* ≤ *b_ij_*. Increasing adaptive knowledge can mitigate the selection cost of a larger brain, but such knowledge is limited by brain size.

Our simulations begin with individuals who have no adaptive knowledge, but the ability to fill their *b_ij_* = 1.0 sized brains with adaptive knowledge through asocial and/or social learning with some probability. To explore the idea that juvenile periods can be extended to lengthen the time permitted for learning, we have included two stages of learning. In both learning stages, the probability of using social learning rather than asocial learning is determined by an evolving *social learning probability* variable (*S_ij_*). We began our simulations with the social learning probability variable set to zero (i.e. at the beginning of the simulation, all individuals are asocial learners). To explore the invasion of asocial learners into a world of social learners, we also ran the simulation with the social learning probability variable set to one (i.e. at the beginning of the simulation, all individuals are social learners). Although social learning is widespread in the animal kingdom (Hoppitt & Laland, 2013), a realistic starting point is closer to pure asocial learning. Nevertheless, the simulations starting with social learners were often useful in understanding these dynamics, so, in some cases, we report these results, as well.

Asocial learning allows for the acquisition of adaptive knowledge, independent of the adaptive knowledge possessed by other individuals. In contrast, social learning allows for vertical acquisition of adaptive knowledge possessed by the genetic parent in the first learning stage or oblique acquisition from more knowledgeable members of the deme (from the parental generation) in the second learning stage. The tendency to learn from models other than the genetic parent is determined by a genetically evolving *oblique learning probability* variable (*V_ij_*). Thus, the simulation does not assume oblique learning or a second stage of learning (a misplaced critique of related models in our opinion; Henrich et al., in press; Vaesen, Collard, Cosgrove, & Roebroeks, 2016; but a critique not relevant to the present model). The probability of engaging in a second round of oblique social learning is a proxy for the length of the juvenile period. In the second stage of learning, if an individual tries to use social learning, but does not use oblique learning, no learning takes place beyond the first stage. This creates an initial advantage for asocial learning and cost for evolution to extend learning into an extended juvenile period. We also allow the ability to select a model with more adaptive knowledge (for oblique learning) to evolve through a *payoff-bias ability* variable (*l_ij_*).

These simulations result in a series of predicted relationships between brain size, group size, adaptive knowledge, asocial/social learning, mating structure, and the juvenile period. Some of these relationships have already been revealed in the empirical literature and thus provide immediate tests of our theory. Specifically, several authors have shown positive relationships (notably in primates) between (1) brain size and social group size (Barton, 1996; Dunbar & Shultz, 2007; Dunbar, 1998), (2) brain size and social learning (Lefebvre, 2013; Reader & Laland, 2002), (3) brain size and length of juvenile period (Charvet & Finlay, 2012; Isler & van Schaik, 2009; Joffe, 1997; Walker, Burger, Wagner, & Von Rueden, 2006), and (4) group size and the length of the juvenile period (Joffe, 1997).

Various hypotheses have been proposed for these relationships. Here we argue that they are all a consequence of a singular evolutionary process, the dynamics of which the CBH models reveal. In addition, we find that different rates of evolutionary change and the size of these relationships across taxa (Shultz & Dunbar, 2010a) may be accounted for by the extent to which adaptive knowledge reduces the death rate (*λ* in our model). As in the analytical model, the *λ* parameter can be interpreted as being part of the resource richness of the ecology. Richer ecologies offer more ‘bang for the buck’, for example, more calories unlocked for less knowledge, allowing individuals to better offset the size of their brains. Higher *λ* suggest a richer ecology. Indeed, research among primates has revealed that factors affecting access to a richer ecology—home range size or the diversity of food sources—are associated with brain size (Clutton-Brock & Harvey, 1980; Harvey & Krebs, 1990). Thus, our model may help explain why both social and ecological variables seem to be variously linked to brain size.

The dynamics of our model also reveal the ecological conditions, social organization and evolved psychology most likely to lead to the realm of cumulative cultural evolution, the pathway to modern humans. These predictions capture the CCBH. Our model indicates the following pathway. Under some conditions, brains will expand to improve asocial learning and thereby create more adaptive knowledge. This pool of adaptive knowledge leads to selection favoring an immense reliance on social learning, with selective oblique transmission, allowing individuals to exploit this pool of growing knowledge. Rogers’ (1988) paradox, whereby social learners benefit from exploiting asocial learners’ knowledge, but do not themselves generate adaptive knowledge, is solved by selective oblique social learning transmitting accidental innovations to the next generation. Under some conditions, an interaction between brain size, adaptive knowledge, and sociality (deme size and interconnectedness) emerges, creating an autocatalytic feedback loop that drives all three—the beginning of cumulative cultural evolution.

#### The Lifecycle

Individuals go through four distinct life stages (see Figure 1): Individuals (1) are born with genetic traits similar to their parents, with some mutation, (2A) learn adaptive knowledge socially from their parents or through asocial learning independent of their parents, (2B) go through a second stage of learning adaptive knowledge through asocial or oblique social learning, (3) migrate between demes, and (4) die or survive to reproduce the next generation. Fecundity and viability selection (birth and death) are expressed separately, allowing us to disentangle the effect of adaptive knowledge on outcompeting conspecifics and on reducing the risk of dying before reproduction.

##### Stage 1: The Birth Stage

In the birth stage, the individuals who survive the selection stage (Stage 4) give birth to the next generation.

**Adaptive Knowledge and the Number of Offspring.** We assume that demes with greater mean adaptive knowledge can sustain a larger population. We formalized this assumption in Equation 13 by linking *k_j_*, which affects the carrying capacity of the deme, to the mean adaptive knowledge of the individuals in the deme (*A_j_*) and some minimum value that we set to our starting group size 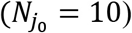. The relationship between mean adaptive knowledge and *k_j_* is scaled by *χ*, but adjusting this coefficient resulted in a computationally intractable deme size as adaptive knowledge accumulated. Therefore, we set this coefficient to a constant value (*χ* = 10) and left exploration of this parameter for a future model. The deme size 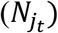 in the current generation (*t*) and *k_j_* are then used to calculate the total expected number of offspring 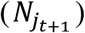 in the next generation (*t* + 1) using the discrete logistic growth function in Equation 14, where *ρ* is the generational growth rate. Initial simulations suggested that *ρ* only affected the rate of evolution rather than the qualitative outcomes. We selected a reasonable value (*ρ* = 0.8) based on Pianka (2011).

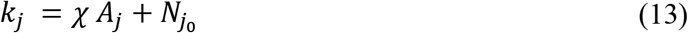

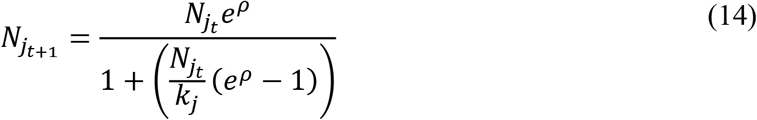

Equation 14 tells us the Expected Value for the number of offspring based on current deme size and *k_j_* (based on deme mean adaptive knowledge). However, this does not tell us which individuals within the deme gave birth to the offspring. We assume that more adaptive knowledge increases an individual’s birth rate. We parameterized the strength of the relationship between adaptive knowledge and birth rate (fecundity selection). A potential parent’s (*i_j_*) probability of giving birth (*p_ij_*) is given by their sigmoid transformed adaptive knowledge value (Equation 15) as a fraction of the sum of all transformed adaptive knowledge values of individuals in the deme (Equation 16). The transformation is adjusted by *φ*, allowing us to study the importance of fecundity selection. For example, we can turn off fecundity selection entirely by setting *φ* = 0: A world with no reproductive skew; all potential parents have the same probability of giving birth. The more we turn up *φ*, the more we have a winner-takes-all world, where to win, one has to acquire adaptive knowledge. This is crucial in thinking about how, for example, our culture-gene co-evolutionary process is influenced by social organization and mating structures that create high reproductive skew.

One mechanism underlying reproductive skew is mating structure. A perfectly monogamous pair-bonded society with no differential selection at the birthing stage would have *φ* = 0. Increasing *φ* allows for an increase in polygyny from “monogamish” (mostly pair-bonded) societies at low values of *φ* to highly polygynous winner-takes-all societies where males with the most adaptive knowledge have significantly more offspring (see Figure 2). Our model suggests that in more polygynous societies, where selection is high, variation is reduced. This allows for the initial rapid evolution of larger brains, but with little or no variation, populations are unable to use social learning to increase their adaptive knowledge and are more likely to go extinct. At the other extreme, evolutionary forces are quashed when *φ* = 0. Social learning and the advent of culture-gene coevolution are more likely to occur when reproductive skew is supressed, such as in monogamish or cooperative/communal breeding societies or where sharing norms result in shared benefits despite skew in ability or success (see Hill & Hurtado, 2009; Hooper, Ross, Mulder, & al., 2016; Lansing et al., 2008). Of course, some argue that culture supports, or is responsible for, such mating structures in humans, which would require us to endogenize *φ*. In our model, we treat *φ* as a parameter.

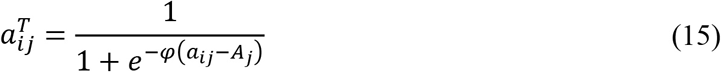

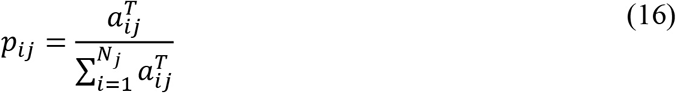

**Figure 2.**
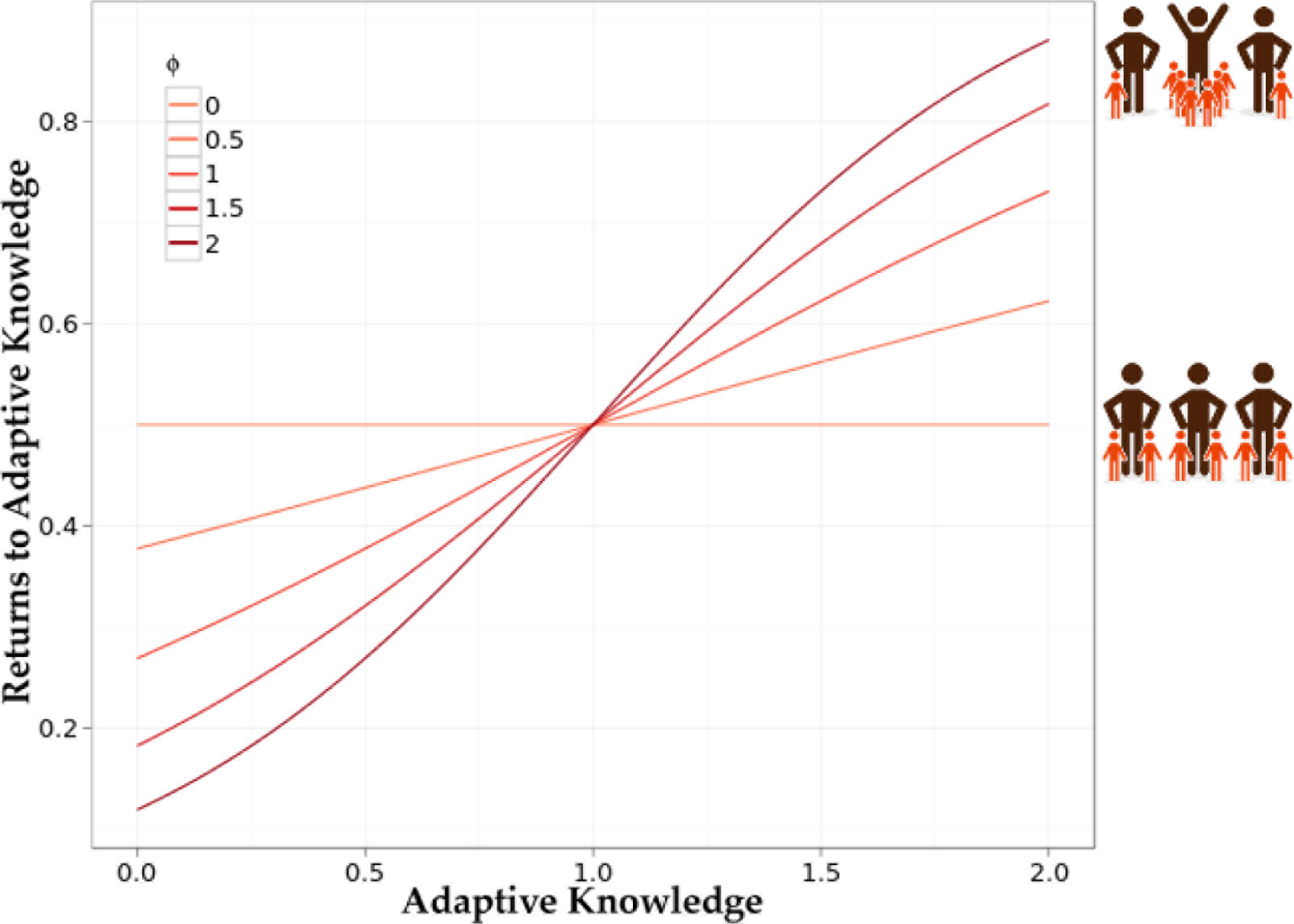
The effect of *φ* on transforming adaptive knowledge. Here the mean adaptive knowledge of the deme is 1 (*A_j_* = 1).

We assume that more individual adaptive knowledge 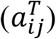 is associated with increased relative fertility. Using a binomial distribution, we instantiate the expected number of offspring *n_ij_* for each parent. A binomial distribution B(*n, p*) describes the number of successes in a sequence of *n* binary experiments (in our model, have offspring vs. don’t have offspring). The probability of success in any particular ‘coin flip’ is given by *p*. For each parent, we draw a value from a binomial distribution where the number of experiments is the Expected Value for the number of offspring in the deme 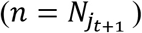 and the probability is calculated by Equation 16, i.e. from 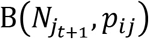. By drawing these values from a binomial distribution, the sum of Expected Values for the offspring of all parents is 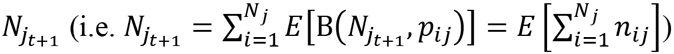.

**Genetic Transmission and Mutation.** The offspring (designated by a prime symbol) born to a parent are endowed with genetic characteristics similar to their parents. These offspring acquire four genetic traits from their parents—their brain size (*b′_ij_*), social learning probability (*s′_ij_*), oblique learning probability (*v′_ij_*), and oblique learning bias (*I′_ij_*). For each trait, newborn individuals have a 1 − *μ* probability of having the same value as their parents (*b_ij_*, *s_ij_*, *v_ij_*, *l_ij_*). If a mutation takes place, new values are drawn from a normal distribution with a mean of their parent value and a standard deviation *σ_s_* for 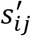, *σ_v_* for *v*′_*ij*_, *σ_l_* for *l*, and *σ_v_* for *v*′_*ij*_ and *σ_b_b_ij_* for 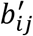. The standard deviations of 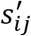 and 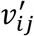 are not scaled by the mean, since these are probabilities and therefore bounded [0,1]. Although 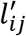 is not bounded, we do not scale the standard deviation by the mean, because small changes in 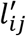 have a large effect on learning bias, due to the sigmoid function. Once offspring have been endowed with genetic characteristics, they then acquire adaptive knowledge. Their method and ability to acquire adaptive knowledge is affected by their genetic traits.

##### Stage 2: Learning

Asocially learned adaptive knowledge values 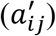 are drawn from a normal distribution based on an individual’s brain size: 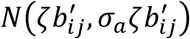. Socially learned adaptive knowledge values are drawn from a similar normal distribution, but with a mean of the model’s (*t*) adaptive knowledge value scaled by transmission fidelity (*τ*): *N*(*τa*_*tj*_,σ_*a*_*τa*_*tj*_). Figure 3 below illustrates the distributions from which these values are drawn and the effect of *ζ* and *τ*.

**Figure 3.**
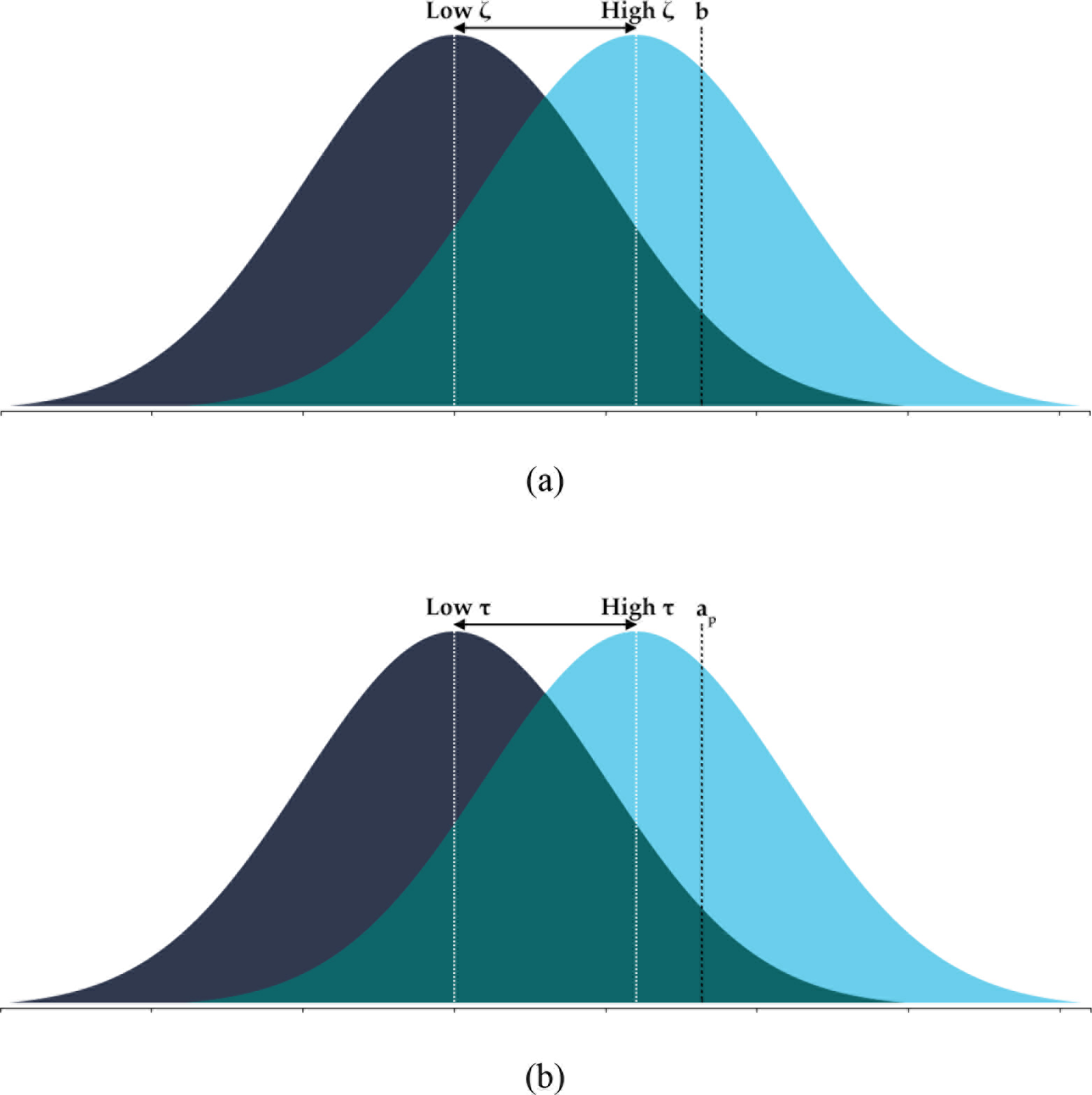
Illustration of distributions for how asocial learning and social learning acquire adaptive knowledge. In (a) an asocial learner has a higher probability of drawing a value closer to their brain size if *ζ* is higher. In (b) a social learner has a higher probability of drawing a value closer to their model’s adaptive knowledge value if *τ* is high. Note that in both cases, adaptive knowledge cannot exceed brain size (*a_ij_* ≤ *b_ij_*).^i^.

For both asocial and social learning, an individual’s adaptive knowledge may not exceed their brain size. But, compared to social learning, asocial learning enables the immediate acquisition of adaptive knowledge based on one’s own brain size. Social learning is dependent on the adaptive knowledge possessed by parents, or those in the parents’ generation within the same deme, if selection extends the learning phrase through a juvenile period.

In Stage 2A, newborn individuals 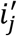 can socially acquire adaptive knowledge from their parent *i* with probability 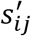. If newborns do not learn from their parents 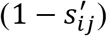, they learn asocially instead.

In Stage 2B, individuals 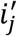 may update their adaptive knowledge through asocial learning with probability 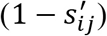 in the same manner as Stage 2A or obliquely from non-parents with probability 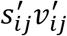. Individuals who do not asocially learn nor obliquely learn do no further learning. This allows us to study conditions under which oblique learning emerges during this extended learning period. Crucially, oblique learning has to out-compete a second round of asocial learning.

We adjust the strength of the relationship between a potential model’s (*m*) adaptive knowledge and their likelihood of being modeled using the learner’s 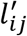 variable in the sigmoid tranformation function (15). A potential model’s (*t_j_*) probability of being selected (*p_tj_*) is given by (16). Notice that these have the same functional form as Equations 15 and 16, and thus the transformation is similar to Figure 2. Both asocial and social learning only update adaptive knowledge values if these values are larger than those acquired during the first stage of learning, Stage 2A.

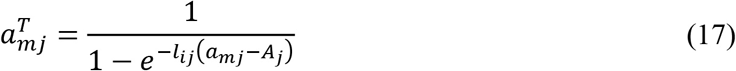

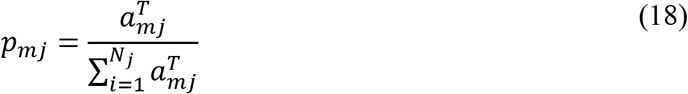

Note, since we are interested in the evolution of social learning, we stacked the deck somewhat against social learning. Individuals have a 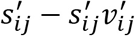 chance of not doing *any* learning during Stage 2B. This creates an initial disadvantage for social learning, since any selection for social learning in Stage 2A risks missing out on a second round of asocial learning in Stage 2B.

##### Stage 3: Migration

Individuals migrate to a randomly chosen deme (not including their own) with probability *m*. All demes have the same probability of immigration. Individuals retain their adaptive knowledge and genetic traits. There is no selection during migration; all individuals survive the journey.

##### Stage 4: Selection Based on Brain Size and Adaptive Knowledge

We formalized the assumption that larger, more complex brains are also more costly using a quadratic function to link brain size to maximum death rate (*c*_*max*_), capturing the idea that the costs of large brains escalate non-linearly with size. In early simulations, we also tested an exponential function, but our exploration revealed no important qualitative differences between the functions.

To formalize the assumption that individuals with more adaptive knowledge are less likely to die *ceteris paribus*, we use the negative exponential function in Equation 19. The *λ* parameter in Equation 19 was varied between simulations and was used to determine the extent to which adaptive knowledge can offset the costs of brain size, where *λ* = 0 indicates no offset. As in our analytical model, the *λ* parameter can be interpreted as how much adaptive knowledge one requires to unleash fitness-enhancing advantages.

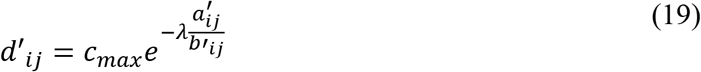

This function captures the idea that the increasing costs of big brains can be offset by more adaptive knowledge. We set *c_max_* = *βb*^2^; *β* = 1/10000 in our simulation). This results in a maximum empty brain size of *b* = 100. The choice of setting the maximum empty brain size to *b* = 100 was somewhat arbitrary, but allowed for a reasonable size brains to see a range of evolutionary behavior (it just sets the scaling). We illustrate the effect of *λ* in Figure 4 below.

**Figure 4.**
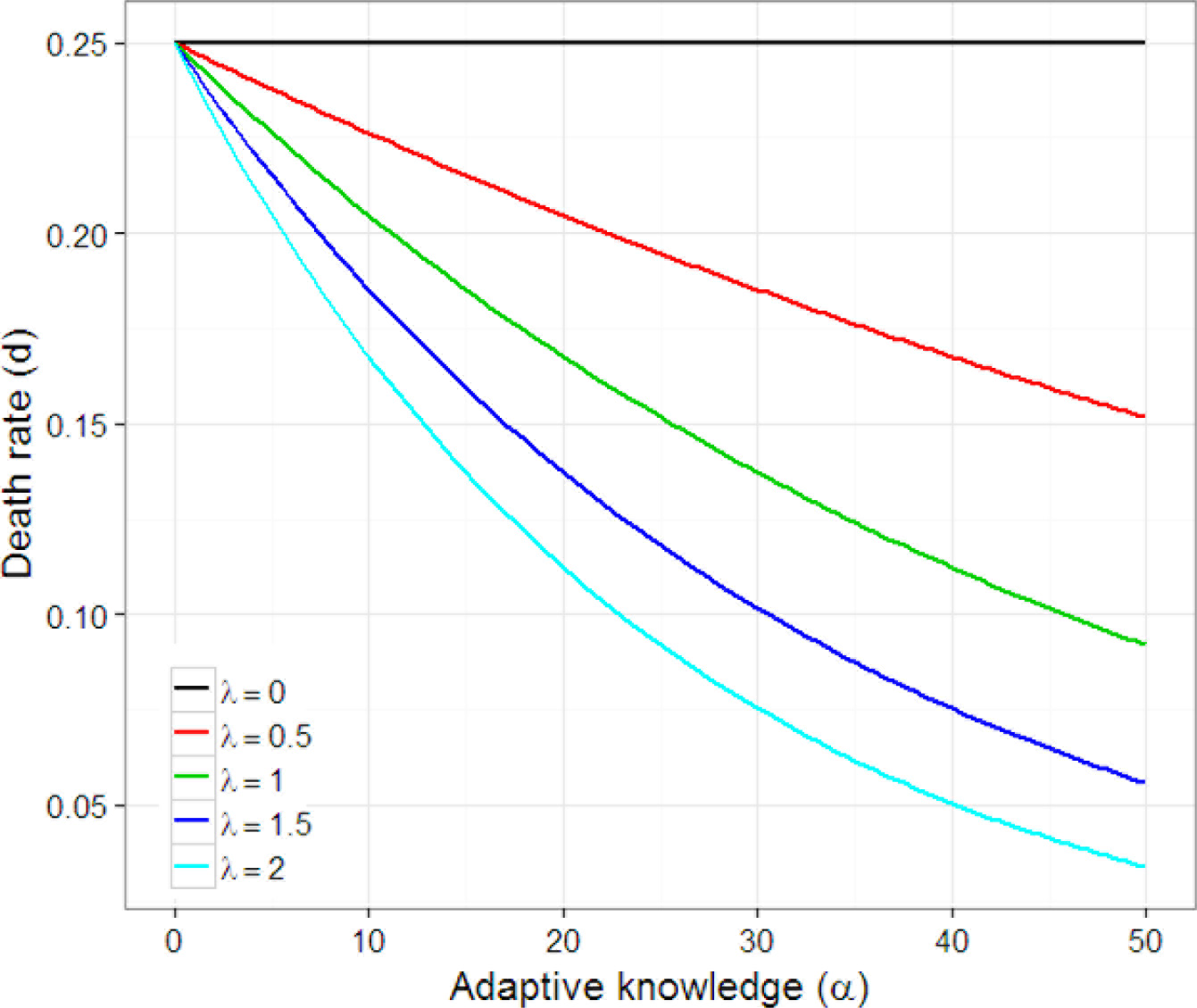
Reduction in death rate for different values of *λ* for a given brain size (*b = 50* in this example).

#### Summary

These basic assumptions generate conflicting selection pressures for (1) more adaptive knowledge and (2) smaller brains. Under some conditions, the cost of having a larger brain is offset by the increased knowledge capacity of larger brains. If adaptive knowledge were freely available, there would be no constraint on the co-evolution of brains and adaptive knowledge; both would ratchet upward. In general, three related constraints prevent this from happening:

1. Adaptive knowledge does not always exist in the environment to fill a larger brain.
2. Larger brains without adaptive knowledge are costly without any offsetting benefits. This is especially true for social learners with brains larger than their parents, since this additional brain space cannot immediately be utilized.
3. Increases in brain size show diminishing returns; brain costs increase at a greater than linear rate.

We simulated a range of space within each parameter set for low, middle, and high values of other parameters for which we found interactions and realistic values of all other parameters. The range for each parameter was as follows: *φ*[0.0,1.0], *τ*[0.75,1.0], *ζ*[0.1,0.9], *m*[0.0,0.2], and *λ*[0.0,2.0].

To give our populations enough time to evolve, we ran our simulation for 200,000 generations. Assuming 25-30 years per generation (Fenner, 2005), this represents 5-6 million years of evolution, approximately the time since the hominin split from chimpanzees (Kumar, Filipski, Swarna, Walker, & Hedges, 2005). With a few exceptions, this guarantees that our genetically evolved variables have hit quasi-equilibrium. To account for stochastic variation in simulation outcomes, we performed 5 iterations per set of unique parameters and averaged the results across these. Unlike the other parameters, learning bias *l* did not generally reach equilibrium; however, we would not expect it to do so since higher *l* values continue to provide an advantage in selecting models, such that *l* should slow down but continue to approach ∞. In our model, *l* is a one-dimensional state variable that captures better and worse ability to select models, but of course in the real world, there are a range of strategies and biases that have evolved to solve the problem of selecting models with more adaptive knowledge. For a discussion of the evolution of these biases and strategies and the trade-offs between them, see Chudek, Muthukrishna, and Henrich (2015) and Henrich (2016). For a list of such biases and strategies, see Rendell et al. (2010).

## Results

We begin by discussing the underlying processes that have led to the relationships between brain size, group size, social learning, and life history observed in the literature. We discuss the effect of our different parameters in creating these relationships and driving evolutionary patterns.

To benchmark the predictions derived by our model, we treat the quasi-equilibrium outcomes of each of our simulation runs as “quasi-species”, with state variables representing the characteristics of each species. We qualitatively compare these simulation outcomes to existing empirical findings in the literature. Then, we focus on the CCBH and examine the conditions that favor substantial amounts of cumulative cultural evolution. The goal here is to understand the conditions under which the interaction between social learning, brain size, group size, sociality and life history generates the kind of auto-catalytic take-off required to explain the last two million years of human evolution.

### The Cultural Brain Hypothesis

Overall, our evolutionary simulations produce patterns that are consistent with the existing empirical data, though, of course, our simulation produces many patterns that have not yet been examined. The causal relationships underlying these patterns—the CBH and our simulated instantiation of it—are outlined in Figure 5 below. Before digging into the details, we summarize these relationships as follows:

1. Larger brains allow for more adaptive knowledge. More adaptive knowledge can, in turn, exert a selection pressure for larger brains.
2. More adaptive knowledge allows for larger potential carrying capacity. Consistent with our analytical model, when there is sufficient adaptive knowledge and transmission fidelity is high enough, there is selection for social learning to take advantage of the adaptive knowledge; larger groups produce more adaptive knowledge that can be exploited by those with better social learning abilities.
3. Large groups of individuals who primarily rely on social learning have larger bodies of knowledge than those who rely on asocial learning, exerting a selection pressure for an extended juvenile period in which more adaptive knowledge can be learned (and created).
4. An extended juvenile period (e.g. adolescence) is a period of reliance on oblique learning (learning from non-genetic parents in the group), which creates a selection pressure for learning biases better able to select individuals and knowledge to learn (better learning abilities and tendency to learn from non-genetic models reinforce each other in a world of plentiful and accumulating adaptive knowledge).
5. Oblique learning and learning biases lead to the realm of cumulative cultural evolution. The length of the juvenile period (period between weaning and sexual maturity) varies across species (Joffe, 1997; Walker et al., 2006), but adolescence (period between sexual maturity and reproduction) may be uniquely human (possible exceptions include elephants (Evans & Harris, 2008) and orca (Olesiuk, Bigg, & Ellis, 1990)). Adolescence may represent a period of oblique social learning, a key to cumulative cultural evolution.

The Cultural Brain Hypothesis predicts that brain size, group size, adaptive knowledge, and the length of juvenile period should be positively intercorrelated among taxa with greater dependence on social learning, but are generally weaker or non-existent among taxa with little social learning. There has been less empirical data published for species with little social learning, perhaps due to a bias toward only publishing statistically significant relationships, making the asocial regime predictions more difficult to test.

The strength of these relationships, overall brain size, and the evolution of different regimes vary, depending on the other parameters in our model. These include ecological factors such as the richness of the ecology (*λ*) as well as other factors that are themselves products of evolution (which we’ve held fixed as phylogenetic constraints): reproductive skew or mating structure (*φ*), transmission fidelity (*τ*), and asocial learning efficacy (*ζ*). Other models have theorized the evolution of these structures, tendencies, and abilities, but here we are interested in the effect of these factors on the co-evolutionary processes shown in Figure 5.

**Figure 5.**
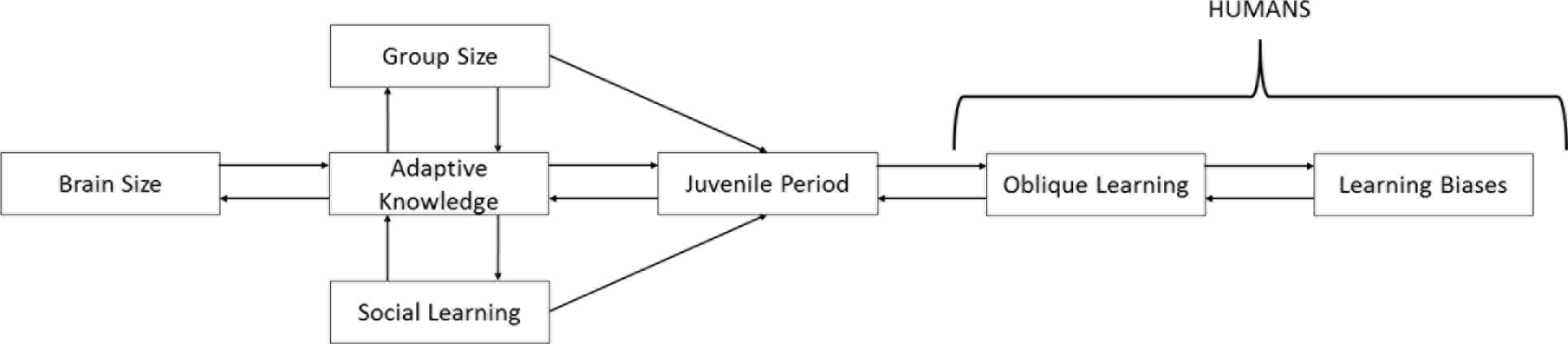
Here we illustrate the causal relationships predicted by the Cultural Brain Hypothesis. Larger brains allow for the storage and management of more information. More adaptive knowledge supports larger brains and larger groups. Larger groups possess more adaptive knowledge for social learning to exploit. Sufficiently large groups of social learners with sufficient knowledge create a selection pressure for a longer juvenile period for social learners to acquire knowledge selectively via biased oblique learning.

#### Effect of Parameters

##### Richness of the Ecology (λ)

Our simulation suggests that the richness of the ecology may be one factor that predicts both the rate of brain evolution and sociality. In a rich ecology (higher *λ*), less adaptive knowledge is needed to unlock more calories, evade more predators, and so on, allowing for larger brains; i.e. adaptive knowledge offers more “bang for the buck”. For those in the realm of social learning, in richer ecologies, we see greater reliance on social learning and larger brains (see Figure 6). Thus the CBH suggests that the empirical correlation that has been shown between sociality and the differential rate of brain expansion between taxa (Shultz & Dunbar, 2010a) may be explained by a third variable: richness of the ecology.

**Figure 6.**
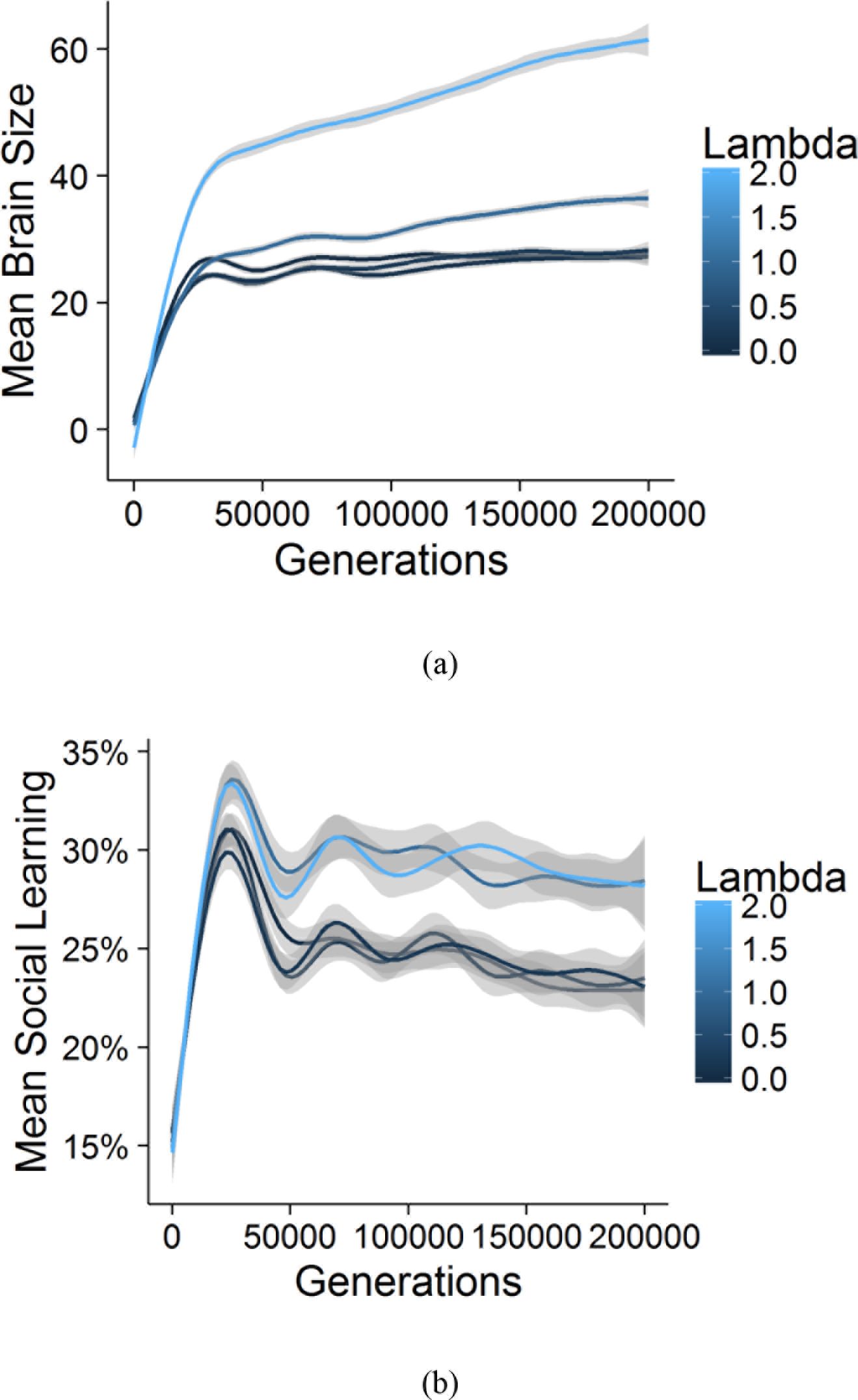
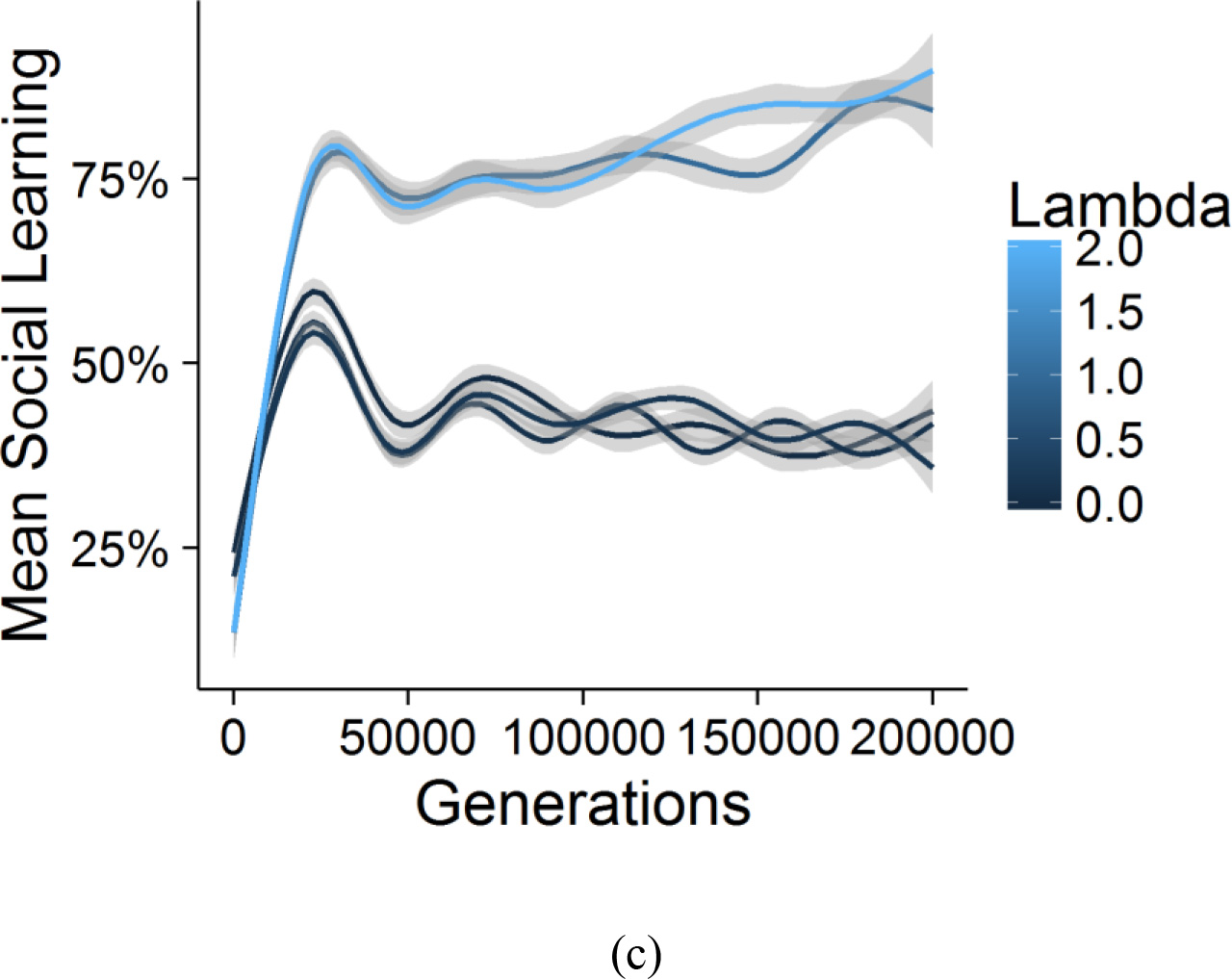
Here we show the effect of richness of the ecology on brain size and social learning. These are aggregated over a range of other parameters (a) Mean brain size showing the encephalization slope for different values of *λ*. Richer ecologies have a steeper slope for brain evolution. (b) Mean social learning showing the slopes over time. Richer ecologies support more social learning when social learning is adaptive. (c) This is made clear in the same plot for a narrower range of other parameters (*τ* = 1 and *ζ* = 0.7).

##### Reproductive Skew or Mating Structure (φ)

We model the effect of mating structure or reproductive skew using *φ*. The *φ* parameter affects the relationship between individual adaptive knowledge and the mating competition. When *φ* = 0, all individuals have the same probability of reproducing regardless of their adaptive knowledge. This corresponds to a perfectly monogamous society with no fecundity selection. As *φ* increases, we enter into a slightly ‘monogamish’ or human cooperative breeding society (where reproductive skew is limited; Hill & Hurtado, 2009) and then to a polygynous society for very high values of *φ*. Increasing *φ*, increases the strength of selection for more adaptive knowledge, but the results of this increase in fecundity selection may be surprising.

First, brain size increases with *φ* (Figure 7a), but this relationship is misleading, because the extinction rate also increases with higher *φ* (Figure 7c). Extinction rates go up, because variance is reduced with too high fecundity selection. More adaptive knowledge is sought at any cost, but in a world with little adaptive knowledge, the best way to acquire this knowledge is via asocial learning. This leads to populations getting stuck in the world of asocial learning without the necessary variance (some attendance and learning from conspecifics) to take advantage of the existing body of adaptive knowledge.

Second, for these same reasons, Figure 7b reveals the tendency to use social learning decreases with greater reproductive skew. We return to this when we discuss the Cumulative Cultural Brain Hypothesis.

**Figure 7.**
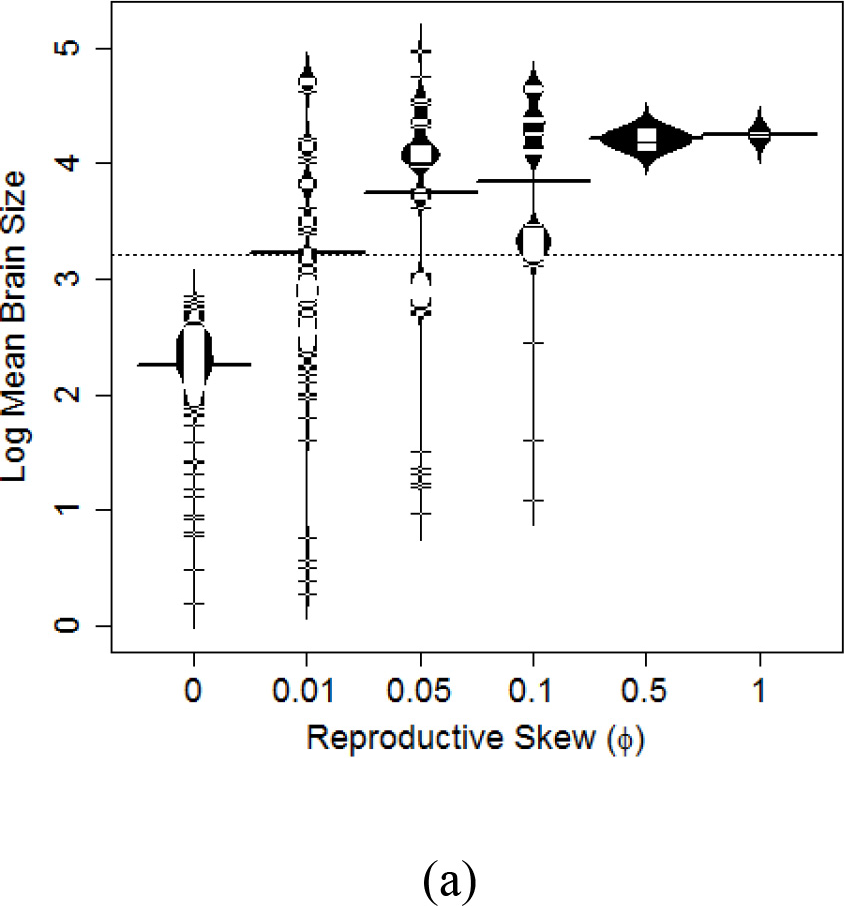
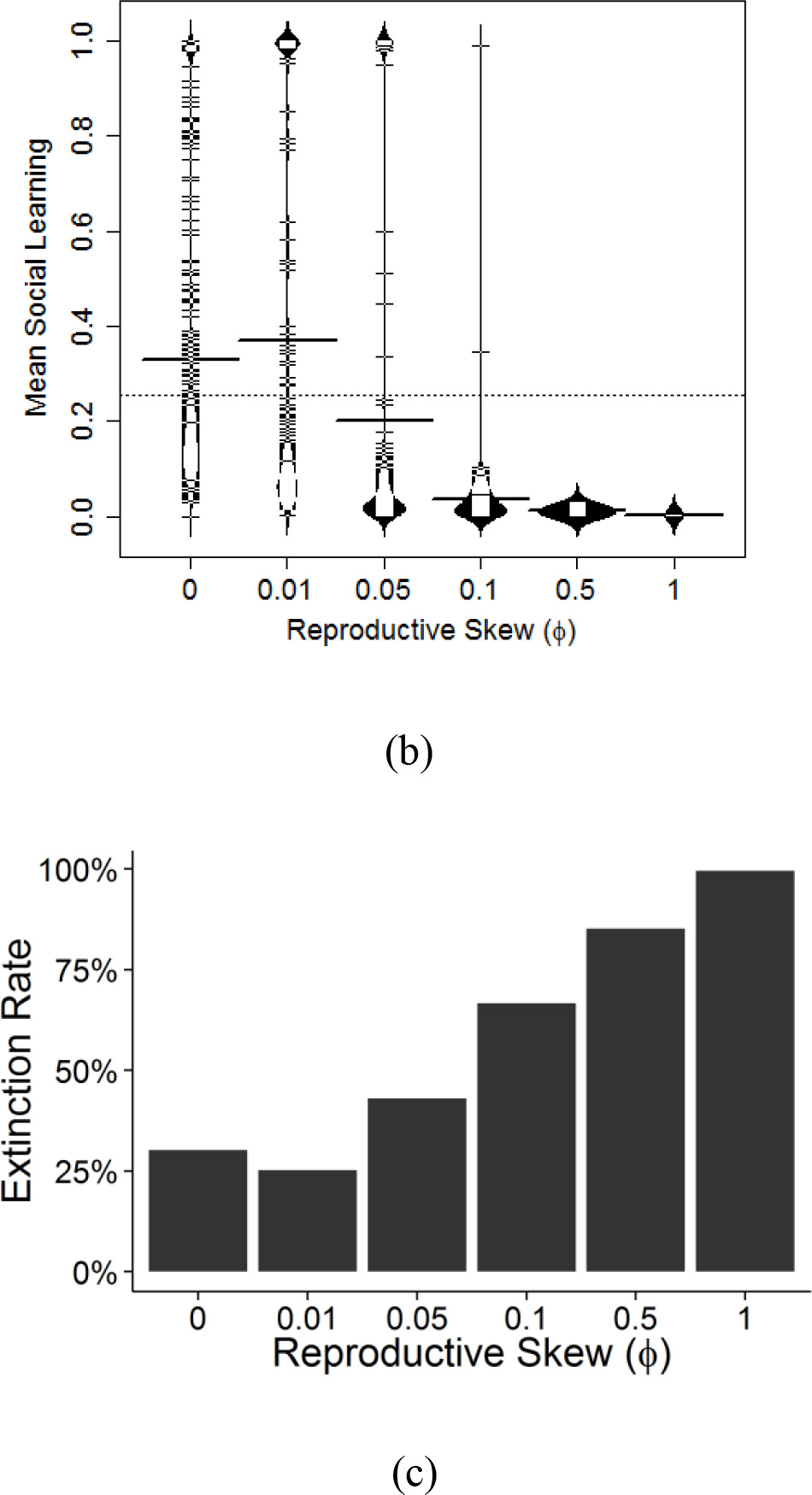
Bean plots showing the distribution of (a) brain size and (b) social learning means for different values of φ. The dotted horizontal line shows the global mean and the bolded horizontal lines show the group means. Bean plots show the distribution of values. (c) Plot showing the rate of extinction for different values of φ.

Empirically, these patterns are consistent with current data: brain size correlates with mating structure in both mammalian and avian lineages (Shultz & Dunbar, 2010a; Shultz & Dunbar, 2010b). Indeed, the relatively high rates of social learning in avian species may be due to their relatively low reproductive skews.

##### Transmission Fidelity and Asocial Learning Efficacy (< and ζ)

Transmission fidelity (*τ*) affects the degree of loss of information in the transmission of adaptive knowledge from cultural models to learners. Asocial learning efficacy (*ζ*) affects the efficiency with which individuals can generate new adaptive knowledge based on their own brain size. In a world of asocial learners, the parameters under which social learning is favored is narrow (recapitulating the insight from Boyd & Richerson, 1996). By starting in a world where the ancestral population has a lot of social learning, we gain two key insights. First, since there is little adaptive knowledge for social learners to take advantage of, we see that asocial learning is initially favored. We discuss this in detail in Section 3 of the Results. Second, with an expanded range in which social learning is favored, we see how *τ* and *ζ* interact in interesting ways to affect the evolution of social learning with consequent effects on brain size, population size, etc. In Figure 8, we plot transmission fidelity against social learning for different levels of asocial learning efficacy where simulations were started with all social learners. Figure 8 shows how social learners can stand on the shoulders of effective asocial learners whose knowledge they exploit. Social learners benefit from smart ancestors.

Although we treat *τ* and *ζ* as parameters in our model, we suspect that if they were allowed to evolve, they would both be pushed higher, as would reliance on social learning. And of course, larger brains that evolve via social learning will also be capable of more potent *asocial* learning since asocial learning is dependent on brain size—both in our model and in reality (see Muthukrishna & Henrich, 2016). We will return to this when we discuss the Cumulative Cultural Brain Hypothesis.

**Figure 8.**
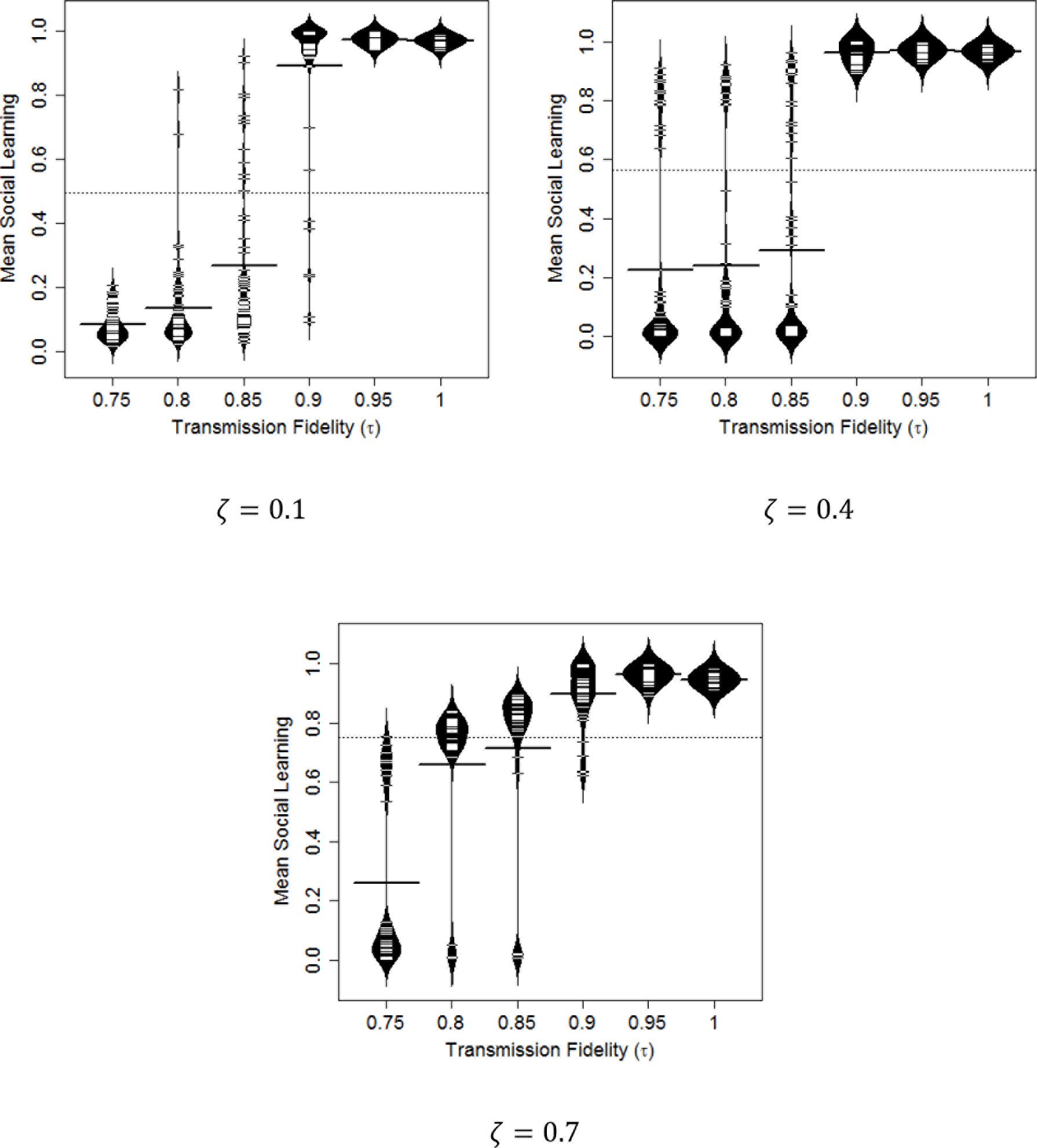
Bean plots showing the distribution of social learning for different values of transmission fidelity (*τ*) and asocial learning efficacy (*ζ*). The dotted horizontal line shows the global mean and the bolded horizontal lines show the group means. Bean plots show the distribution of values. Transmission fidelity interacts with asocial learning efficacy to generate high equilibrium reliance on social learning.

To most effectively compare our theoretical findings to the existing empirical data, we subject our simulation output to the same kinds of analyses used by researchers in the empirical literature. Of course, this comparison is qualitative: we didn’t select parameter values to fit the empirical literature, but instead sought to use a wide range of plausibly realistic values, so we don’t expect exact matches between the empirical correlations and our theoretical predictions. There’s little doubt that some of our parameter setting never or rarely occur in the real world.

#### Predictions

Our range of parameters results in a range of simulated quasi-species (referred to as “species” from herein) with predicted relationships between the characteristics of these species. We have 4 key parameters in our model: Reproductive skew (mating structure; *φ*), transmission fidelity (*τ*), asocial learning efficacy (*ζ*), and richness of the ecology (*λ*). Each represents different ecological and phylogenetic constraints. The species that emerge under different combinations of these conditions can be partitioned into at least two regimes (Figure 9): species that mostly rely on (1) asocial learning or (2) social learning. A k-cluster analysis on the mean social learning value (*s*) for each simulation run suggests that the threshold between these regimes is approximately 50%. Note that the relative count size of the two regimes is a reflection of the range of parameters we chose rather than a reflection of the world (e.g., transmission fidelity values greater than 75%, rather than from 0% to 100%). Under some conditions, a species that mainly relies on social learning can enter into the realm of cumulative cultural evolution. The conditions that predict this transition are the basis of the CCBH. The relationships between equilibrium state variable values differ considerably between these two regimes and so we analyze them separately. The species that mostly rely on social learning include those in the realm of cumulative cultural evolution.

**Figure 9.**
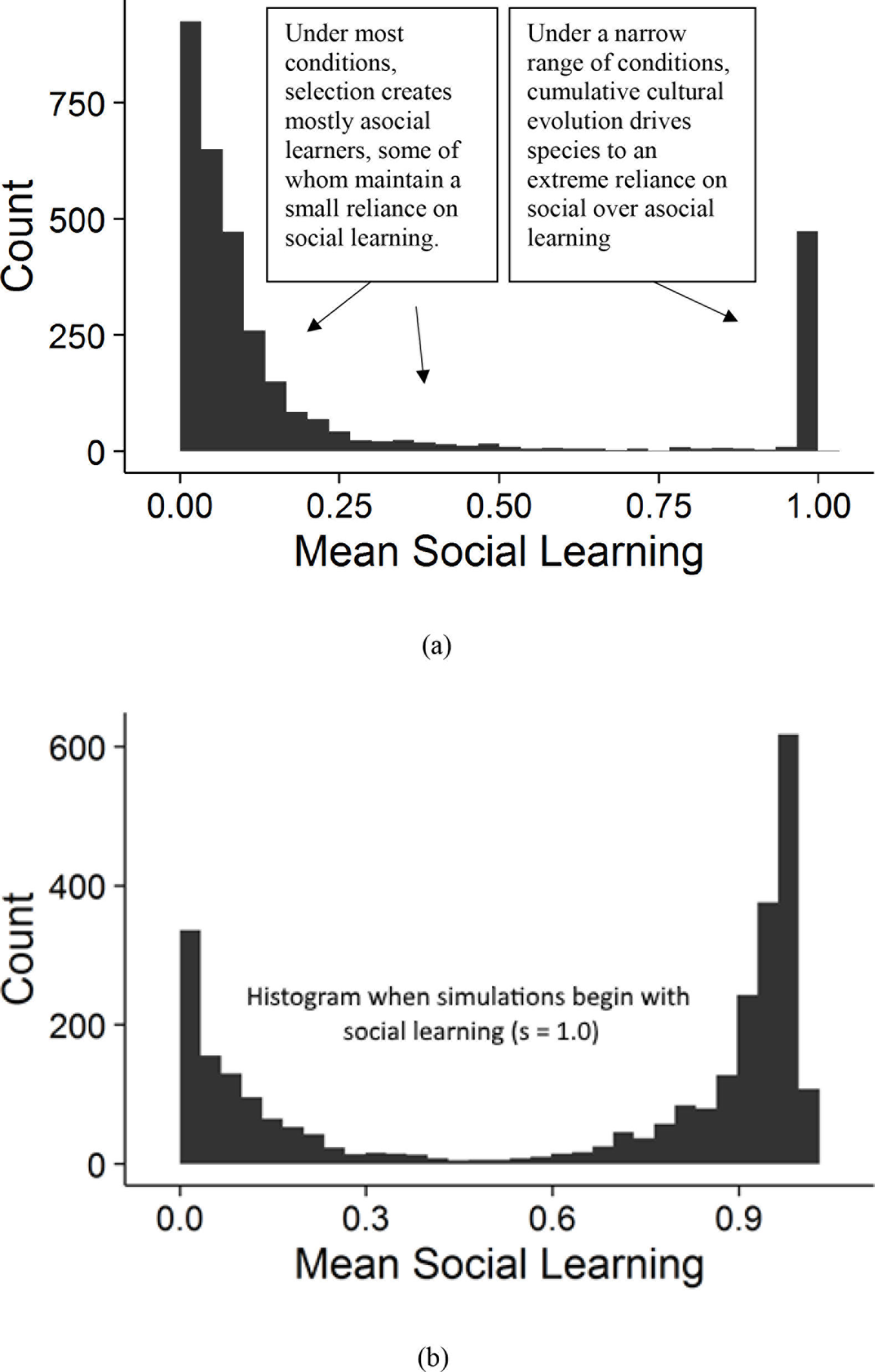
Histogram of mean social learning probability (*s*). Under most conditions, selection creates individuals primarily reliant on asocial learning, some of whom maintain a small reliance on social learning. Under a narrow range of conditions, cumulative cultural evolution drives species to an extreme reliance on social over asocial learning. Consistent with previous models (e.g. Boyd & Richerson, 1996), this range of conditions expands if social learning is assumed to exist in the ancestral species; i.e., if we start the simulation with social learners. (b) Histogram of mean social learning probability (*s*) when simulations began with all social learners (*s* = 1.0).

To confirm that the relationships we report are not driven by cumulative cultural species (humans or hominins), we also ran a k-cluster analysis assuming 3 regimes. This analysis split species into primarily asocial learners (*s* < 0.20; e.g., beetles and buffalo), a few species with some reliance on social learning (0.20 < *s* < 0.66; e.g., capuchins and chimpanzees), and species that are almost entirely reliant on social learning (*s* > 0.66; e.g., humans, hominins, and close cousins). We then show that the relationships we find among species that mainly rely on asocial learning (*s* < 0.50) also hold among highly asocial learning species (*s* < 0.20), and relationships we find among species that mainly rely on social learning (*s* > 0.50) also hold among species with some social learning (0.20 < *s* < 0.66).

#### Testing Predictions

We can test our theoretically-derived qualitative predictions by comparing the species that emerge in our simulation with empirical data. Table 2 reports the relationships between the evolved characteristics of our species for each regime in our range of parameters. Below, we feature 4 key predicted relationships—(1) brain size vs. group size, (2) brain size vs. social learning, (3) brain size vs. juvenile period, and (4) group size vs. juvenile period.

**Table 2.** Correlations for each regime across our entire parameter space. Correlations between log mean brain size, log mean adaptive knowledge, log mean group size, mean social learning, and mean juvenile period with 95% confidence intervals in brackets. The table has been color coded from red (*r* = −1) to white (*r* = 0) to blue (*r* = 1) for ease of comprehension. The upper table has correlations across the entire parameter space. The lower table has primarily asocial learners (*s* < .5) in the bottom triangle and primarily social learners (*s* > .5) in the top triangle. Following the empirical literature, social learning is defined as the number of observed incidents of social learning. Thus, we multiplied *s* by mean group size (*N*), and then following the empirical work, added 3, and took the natural log (Reader & Laland, 2002). The juvenile period is defined as the probability of socially learning in a second round of learning (*sv*). Higher *sv* values should demand a longer juvenile period.

| Evolving characters | | Brains<br>$\log(\bar{b})$ | Adaptive Knowledge<br>$\log(\bar{a})$ | Group Size<br>$\log(\bar{N})$ | Social Learning<br>$\log(3 + \bar{N}s)$ | Juvenile Period<br>$\bar{sv}$ |
| --- | --- | --- | --- | --- | --- | --- |
| Across entire parameter space | $\log(\bar{b})$ | 1 | | | | |
| | $\log(\bar{a})$ | 0.81 [.78,.82] | 1 | | | |
| | $\log(\bar{N})$ | 0.51 [.48,.53] | 0.62 [.60,.64] | 1 | | |
| | $\log(3 + \bar{N}s)$ | 0.17 [.14,.20] | 0.42 [.39,.45] | 0.64 [.62,.66] | 1 | |
| | $\bar{sv}$ | 0.05 [.01,.08] | 0.24 [.21,.27] | 0.27 [.24,.30] | 0.81 [.80,.82] | 1 |
| Primarily asocial: bottom | $\log(\bar{b})$ | 1 | 0.99 [.98,.99] | 0.72 [.67,.75] | 0.72 [.68,.76] | 0.17 [.09,.25] |
| | $\log(\bar{a})$ | 0.78 [.77,.80] | 1 | 0.69 [.65,.74] | 0.70 [.66,.74] | 0.15 [.06,.23] |
| | $\log(\bar{N})$ | 0.42 [.39,.45] | 0.55 [.53,.58] | 1 | 0.98 [.98,.98] | 0.22 [.14,.30] |
| | $\log(3 + \bar{N}s)$ | -0.23 [-.26,-.19] | 0.13 [.10,.17] | 0.61 [.58,.63] | 1 | 0.22 [.14,.30] |
| | $\bar{sv}$ | -0.53 [-.56,-.51] | -0.34 [-.37,-.30] | -0.21 [-.25,-.18] | 0.42 [.39,.45] | 1 |

##### Brain Size and Group Size

As Table 2 shows, our model indicates that among species that *mainly rely on social learning* (defined as *s* > 0.5), the relationship between brain size and group size is *r* = 0.72 [0.68,0.76]. Among species with some social learning (0.20 < *s* < 0.66), the correlation is similarly, *r* = 0.72 [0.66,0.77]. In contrast, our model predicts that among taxa that rely more on asocial learning, the relationship is much weaker, *r* = 0.42 [0.39,0.45]. Among highly asocial learners (*s* < 0.20), the correlation is *r* = 0.35 [0.30,0.37].

The empirical literature has established a strong positive relationship between brain size and group size in primates, but not in other taxa (Dunbar, 2009; Fox et al., 2017; Pérez-Barbería, Shultz, & Dunbar, 2007; Shultz & Dunbar, 2007). In primates, the correlation between relative neocortex size and group size is somewhere between *r* = 0.48 to *r* = 0.61 (Barton, 1996). We contrast our theoretical predictions to the empirical data in Figure 10. In support of the SBH, researchers have noted that in other taxa, brain size correlates with measures of sociality or social group complexity (e.g., among non-primate mammals Fox et al., 2017; Shultz & Dunbar, 2010a; Shultz & Dunbar, 2007) and with mating structure (e.g. among birds; Shultz & Dunbar, 2010b). However, why group size correlates with brain size in some taxa and not others remains a mystery (Dunbar & Shultz, 2007). The CBH offers an explanation, predicting that the strength of the relationship between brain size and group size increases with reliance on social learning due to larger groups offering a greater number of opportunities for social learning and a greater amount of information to learn. Thus, for example, we should expect (and do see) a relationship between brain size and group size in primates, but not ungulates or carnivores (who display less social learning; van Schaik & Burkart, 2011).

**Figure 10.**
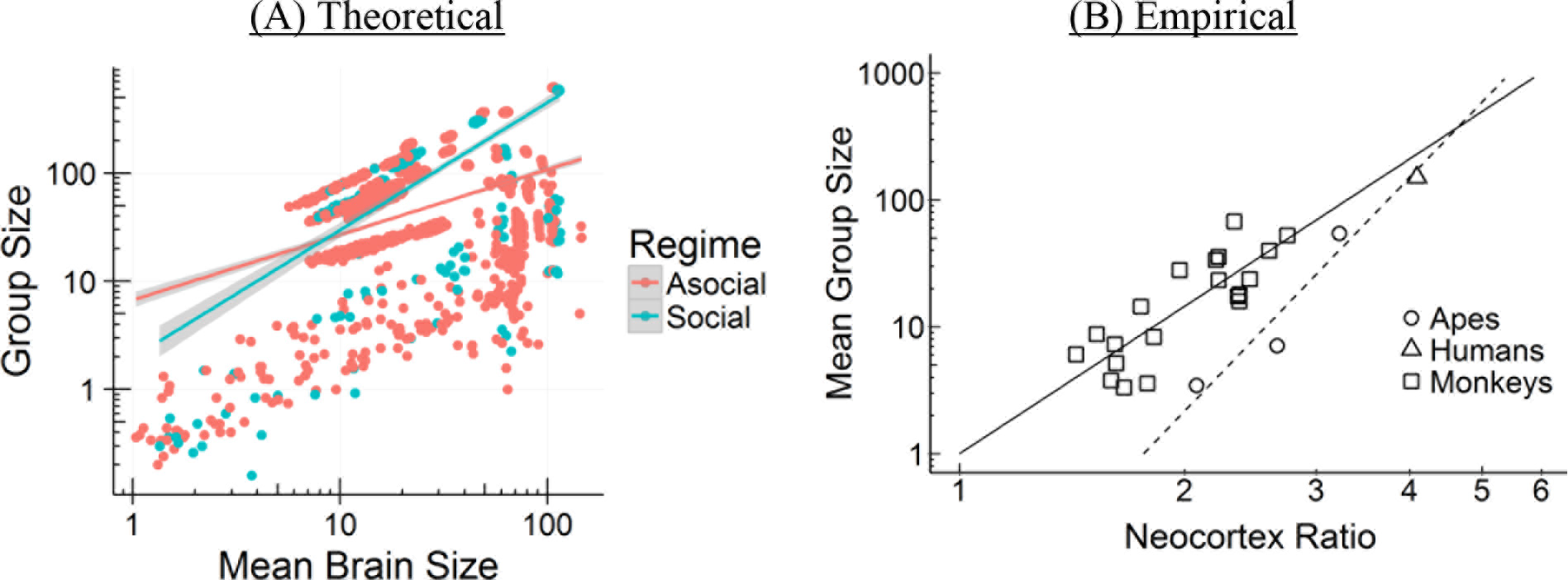
Brain size and group size. (A) Our model’s empirical correlations between brain size and group size (*r* = 0.42 [Asocial], *r* = 0.72 [Social]). (B) Empirical correlation between brain size and group size from Barton (1996) is somewhere between *r* = 0.48 to *r* = 0.61.

Notably, the algorithms in our theoretical model do not specify a direct relationship between brain size and group size or group size and brain size—these relationships just emerge. Instead, the CBH assumes that larger brains are better at storing and managing adaptive knowledge. There are two pathways to acquire that knowledge: asocial learning and social learning. Groups with higher mean adaptive knowledge have a higher carrying capacity, thus taxa more reliant on asocial learning generally also have a positive relationship between brain size and group size in our model. For taxa more reliant on social learning, larger groups also have more adaptive knowledge to exploit, raising the mean adaptive knowledge of the group and therefore the carrying capacity. Thus, our model predicts a stronger relationship between brain size and group size among taxa more reliant on social learning (compared to those more reliant on asocial learning).

##### Brain Size and Social Learning

Our simulations reveal a positive relationship between brain size and social learning across species. Among species that primarily rely on social learning (*s* > 0.5), the relationship between brain size and social learning is *r* = 0.72 [0.68,0.76]. Among species with some social learning (0.20 < *s* < 0.66), the correlation is *r* = 0.58 [0.49,0.65]. However, among species that primarily rely on asocial learning (*s* < 0.5), the relationship is negative, *r* = −0.23 [−0.26,−0.19] and similarly in strongly asocial learning species, (*s* < 0.20): *r* = −0.24 [−0.27,−0.20]. Most asocial learning species remain small brained, but those that do acquire larger brains via genetically-hardwired asocial learning do so at the expense of much reliance on social learning abilities.

It bears emphasis that the trade-off here is between *time* or *effort* spent on asocial vs. social learning, not between brain tissue allocation. If you are doing asocial learning—say running trial and error to improve a tool—you can’t be carefully watching others at the same time. Or, alternatively, sometimes the suggested behavior delivered by asocial vs. social learning processes will be contradictory, and organisms have to decide which source they will rely on. In both of these senses, there’s an unavoidable trade-off between social and asocial learning. However, in our model, bigger brains are always better at asocial learning (when they do it), even if the selection pressure that drove that brain expansion was due to the effects of social learning. That is, we assume complementarity as suggested by Reader et al. (2011); and Reader and Laland (2002).

From the empirical literature, social learning is measured by observational counts of social learning events, and reveals a correlation with brain size of *r* = 0.69, *p* < 0.001 (*r* = 0.36, *p* < 0.05, controlling for phylogeny) for primates (Lefebvre, 2013; Reader & Laland, 2002). To better match our social learning probability, *s*, to the empirically available results, we assumed that simulated species with larger populations and higher *s* values would generate greater numbers of observational counts (linearly). Thus, we multiplied *s* by mean group size (*N*), and then following the existing empirical approach, added 3, and took the natural log (Reader & Laland, 2002). A similar relationship has been shown for birds using indirect measures of opportunities for social learning (e.g. number of caretakers; van Schaik et al., 2012). Figure 11 contrasts our predicted relationship with the empirical literature.

**Figure 11.**
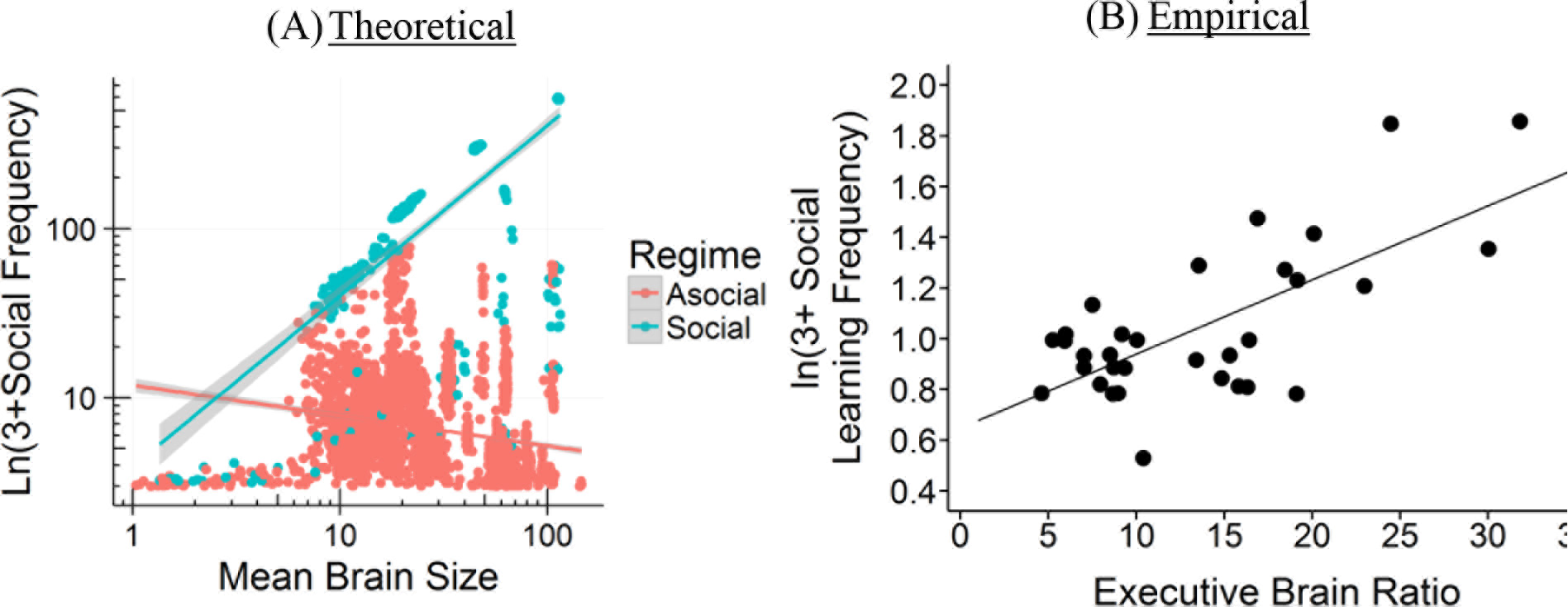
Brain size and social learning. (A) Our model’s empirical correlations between brain size and incidences of social learning (*r* = −0.23 [Asocial], *r* = 0.72 [Social]). (B) Empirical correlation between brain size and incidences of social learning among primates from Reader and Laland (2002) is *r* = 0. 69 (*r* = 0. 36 controlling for phylogeny). A similar relationship has been shown for birds using indirect measures of opportunities for social learning (e.g. number of caretakers; van Schaik et al., 2012).

##### Brain Size and Juvenile Period

Our simulation does not explicitly model the length of the extended juvenile period, but does include 2 periods of learning. In the first period, individuals can learn socially from their genetic parent or asocially by themselves. In the second period, individuals with a low s value are likely to update their knowledge asocially, while those with higher *s* values only updated their knowledge obliquely based on their *v* value; individuals had a 1 − *s* probability of updating asocially, *sv* probability of updating socially and an *s* − *sv* probability of doing no further learning. Thus, *sv* represents an extended juvenile period in which learners could use payoff-biased oblique transmission to update their knowledge. Larger *sv* values should demand a longer juvenile period.

Our model indicates that among species that mainly rely on social learning (*s* > 0.5), the relationship between brain size and the length of the extended juvenile period is *r* = 0.17 [0.09,0.25]. This positive relationship only occurs when we include highly social learners (*s* > 0.66). The relationship between brain size and the length of an extended juvenile period disappears or is negative among species with only a moderate reliance on social learning (0.20 < *s* < 0.66), *r* = −0.08 [−0.20, 0.04], more reliant on asocial learning (*s* < 0.5), *r* = −0.53 [−0.56,−0.51], or are highly asocial (*s* < 0.20), *r* = −0.59 [−0.61,−0.56]. Thus, we argue that an extended juvenile period evolves to support more opportunities to engage in social learning.

Our *extended* juvenile period most closely represents an adolescent period (the period from sexual maturity to sexual reproduction), where additional biased oblique social learning occurs. Adolescence is rare, occurring in humans, possibly elephants (Evans & Harris, 2008) and some orca (Olesiuk et al., 1990), and some members of cooperative breeding species (Hawn, Radford, & du Plessis, 2007). Nonetheless, positive relationships between brain size and the length of the *juvenile* period (weaning age to sexual reproduction) have been shown directly in primates (Charvet & Finlay, 2012; Joffe, 1997; Walker et al., 2006) and indirectly via age to sexual maturity in a variety of taxa (Isler & van Schaik, 2009). The correlation for primates is *r* = 0.61, *p* = 0.037 (Joffe, 1997). Though the comparison is imperfect, we show the relationship between brain size and length of the extended juvenile period side by side with the relationship between brain size and the juvenile period in primates in Figure 12 below.

**Figure 12.**
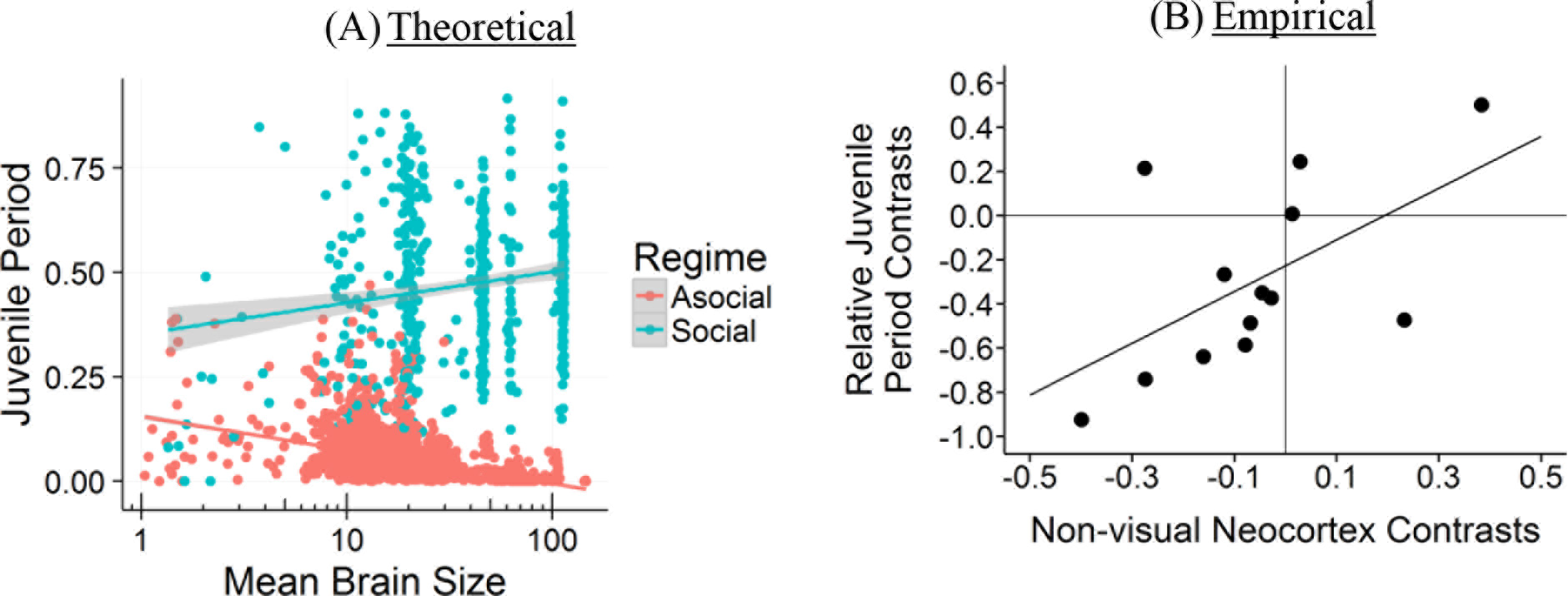
Brain size and the juvenile period. (A) Our model’s empirical correlations between brain size and the length of the extended juvenile period (*r* = −0.53 [Asocial], *r* = 0.17 [Social]). (B) Empirical correlation between brain size and juvenile period among primate species from Joffe (1997) is *r* = 0.61.

##### Group Size and Juvenile Period

Since an extended juvenile period primarily evolves in the presence of large amounts of adaptive knowledge that requires more opportunities for social learning, we should also expect to see a positive relationship between group size and the juvenile period among highly social learners. Indeed, our model indicates that among species that mainly rely on social learning (*s* > 0.5), the relationship between group size and the length of the juvenile period is *r* = 0.22 [0.14,0.30]. As with the relationship between brain size and the length of the extended juvenile period (and for related reasons), this positive relationship only occurs when we include highly social learners (*s* > 0.66). The relationship between brain size and the length of an extended juvenile period disappears or is negative among species with only a moderate reliance on social learning (0.20 < *s* < 0.66), *r* = −0.05 [−0.16, 0.07], mainly rely on asocial learning (*s* < 0.5), *r* = −0.21 [−0.25, −0.18], or are highly reliant on asocial learning (*s* < 0.20), *r* = −0.12 [−0.16, −0.08]. For highly social learning species, this positive relationship is an indirect consequence of social learners having access to more knowledge in larger groups, creating a stronger selection pressure for a longer juvenile period in which to take advantage of this knowledge. This, in turn, raises the average adaptive knowledge of the group, allowing for larger groups. Empirically, in primates, the relationship between absolute juvenile period length (we were unable to find the weaning age to sexual maturity measure; sexual maturity to sexual reproduction is non-existent) and mean group size is *r* = 0.57, *p* = 0.007 (Joffe, 1997). In Figure 13 below, we contrast our predictions against the empirical results. Joffe (1997) did not provide a comparison plot, but we have generated one from his data.

**Figure 13.**
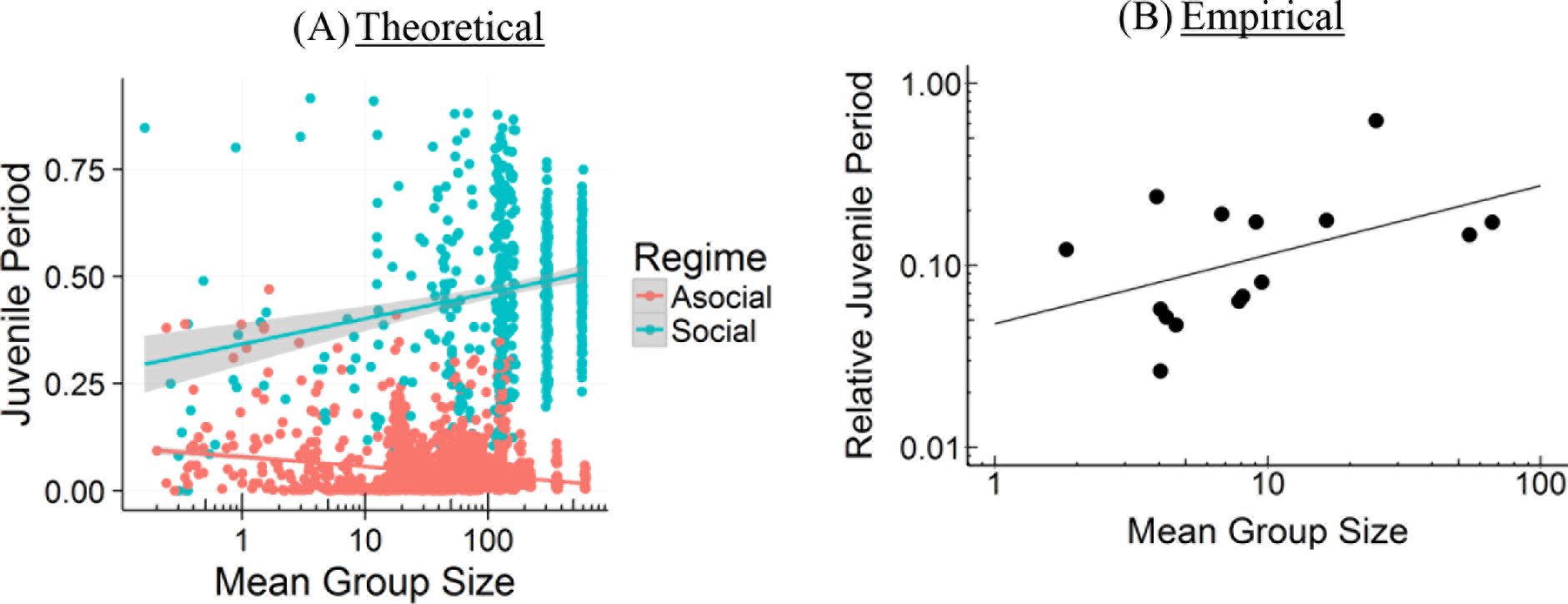
Group size and the juvenile period. (A) Our model’s empirical correlations between group size and the length of the juvenile period (*r* = −0.21 [Asocial], *r* = 0.22 [Social]). (B) Empirical correlation between group size and the length of the juvenile period among primates from Joffe (1997) is *r* = 0. 57.

### The Cumulative Cultural Brain Hypothesis

Beyond the hypothesis that social learning, brain size, adaptive knowledge, and group size may have coevolved so as to create the patterns found in the empirical literature, we are also interested in the conditions under which these variables might interact synergistically to create highly social species with large brains and substantial accumulations of adaptive knowledge (humans). To assess when an accumulation of adaptive knowledge becomes cumulative cultural evolution, we apply a standard definition of cumulative cultural products as being those products that a single individual could not invent by themselves in their lifetime. To calculate this for our species, we ask what the probability is that an individual with the average brain size of the species would invent the mean level of adaptive knowledge in that species via asocial learning.

#### Formalization of Cumulative Culture

The probability of an individual *i* in deme *j* acquiring the mean deme adaptive knowledge *A_j_* through asocial learning is given by Equation 20. Asocial learners draw their adaptive knowledge value from a normal distribution with mean of their brain size scaled by *ζ*. Thus the probability of acquiring *a_ij_* ≥ *A_j_* is the integral from this mean value or greater over the asocial learning distribution. Note that this gives the probability of an individual acquiring that level of adaptive knowledge. The probability that the mean adaptive knowledge of the deme is reached through asocial learning is this probability to the power of the number of individuals in the deme 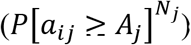—a slim chance indeed.

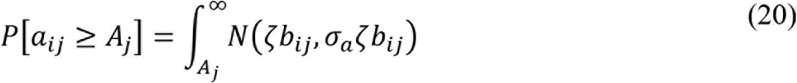

We set a low, albeit arbitrary, threshold where the probability of any individual acquiring this level of adaptive knowledge through asocial learning is less than 0.1%. At this level, the probability that an entire population would develop that level of adaptive knowledge through asocial learning is 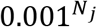, i.e., exceedingly unlikely. Thus, mean levels of adaptive knowledge that are so exceedingly unlikely to have been acquired through asocial learning can be attributed to cumulative cultural evolution. In Figure 14 below, we plot brain size against the probability of acquiring that amount of information.

**Figure 14.**
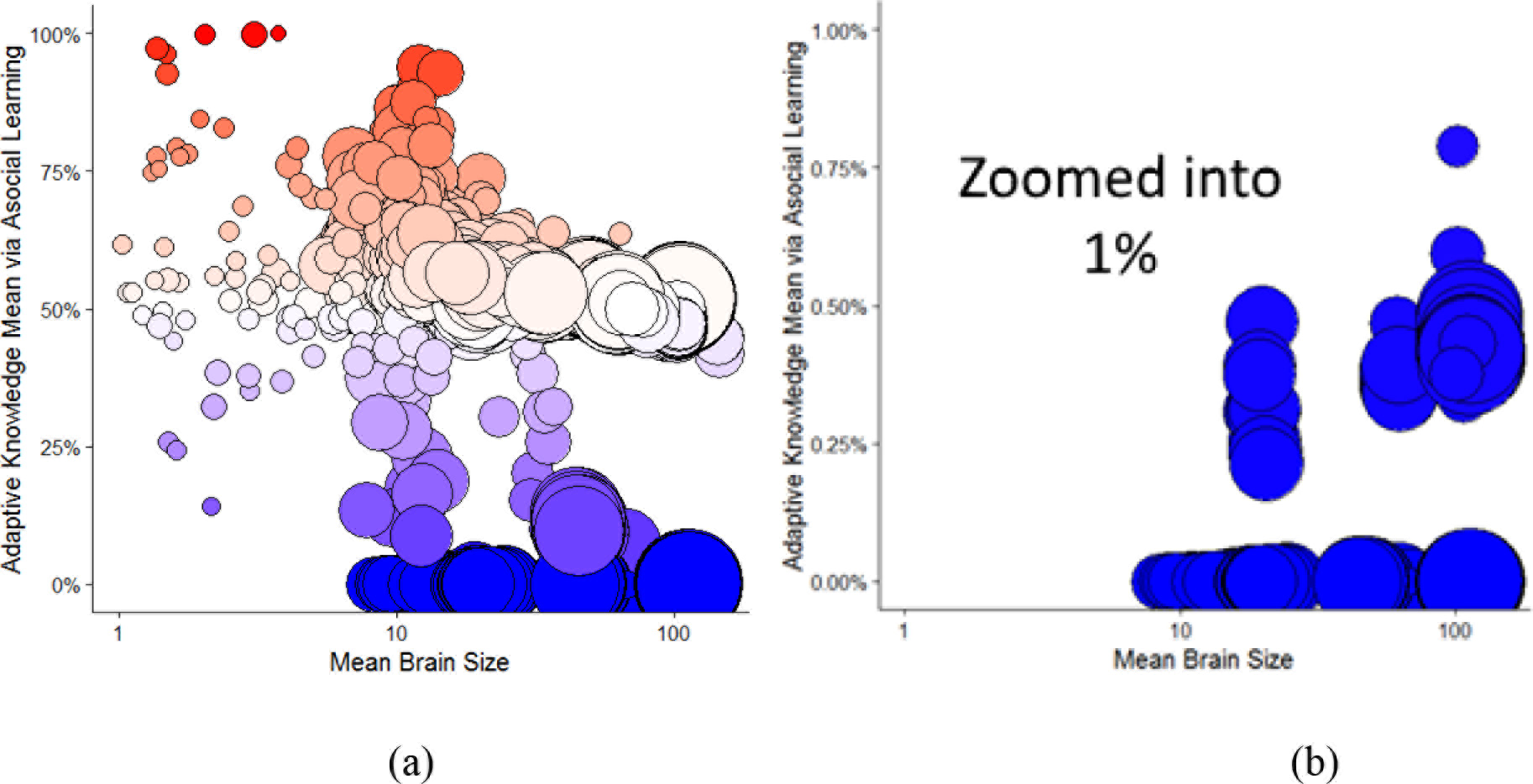
Cumulative Culture and Brain Size. Circle size indicates the mean population size. More red indicates high probability of acquiring knowledge through asocial learning and more blue indicates a low probability. The darkest blue circles in the bottom right are the simulations that cross the threshold into the cumulative cultural realm. (a) Log mean brain size against the probability of acquiring the mean adaptive knowledge in the group via asocial learning. (b) Here we show the same data zoomed in-between 0 and 1%.

Next, we can look at what parameters increase and decrease the probability of entering into the realm of cumulative cultural evolution, on the bottom right corner of Figure 14a: those largebrained species with a lot of adaptive knowledge, which they were unlikely to acquire without cumulative cultural evolution.

#### Transmission Fidelity Drives Larger Brains

The simulation predicts that transmission fidelity is the key to entering into the realm of cumulative cultural evolution. When the model begins with widespread social learning, we see a threshold effect, where for very high fidelity transmission (*τ* > 0.85), social learning and large brains evolve under a wide range of parameters. However, when we begin with primarily asocial learners (more plausible), this threshold increases to nearly 100% (see Figure 16). The degree of these results may be exaggerated by our “stacking the deck” against social learning, but the overall results are consistent with previous models that argue that transmission fidelity is the key to cumulative cultural evolution (Lewis & Laland, 2012). And also with models that show that there is a fitness valley that needs to be crossed to enter into the realm of cumulative culture and reliance on social learning (Boyd & Richerson, 1996). When social learning is already present in the population, species can enter the realm of cumulative cultural evolution under a wider range of parameters—that is, the more pre-existing social learning exists, the shallower the fitness valley that needs crossing.

Embedded in *τ*, and eventually oblique learning and learning bias, are cognitive abilities like theory of mind, the ability to recognize, distinguish, and imitate potential models, but also teaching and social tolerance. We suspect that if we endogenized *τ*, either cultural or genetic evolution would favour higher values under these conditions.

#### Mating Structure Matters

As discussed, lower reproductive skew consistent with “monogamish” or human cooperative/communal mating structures (Hill & Hurtado, 2009) are more likely to lead to social learning and therefore to cumulative cultural evolution. Too strong a selection pressure leads to bigger brains via asocial learning—bigger mutant brains can’t be filled via social learning, since cultural information is capped by brain size—but these populations often go extinct, even when we start with fully developed social learning. We graph the probability of entering into the realm of cumulative cultural evolution for different values of *φ* in Figure 16a. We see a Goldilocks’ zone around *φ* = 0.01, regardless of the starting conditions (though as previously discussed, the parameter range leading to cumulative culture increases if social learning is common). As reproductive skew (e.g., polygyny) increases, asocial learning is favored. Thus, entering the realm of cumulative cultural evolution is less likely.

#### Smart Ancestors and Rich Ecologies

As discussed in (1), we find that an interaction between transmission fidelity *τ* and individual learning *ζ* fuels the autocatalytic take-off. If *ζ* is too high, individual learning is too efficient and social learning struggles to take flight, except at very high rates of transmission fidelity or if social learning is already present. But if *ζ* is too low, even if social learning out- competes individual learning, populations have smaller brains and less adaptive knowledge compared to when social learning out-competes more effective individual learning. These results suggest that social learners stand on the shoulders of effective asocial learners. That is, when social learning can initially exploit the adaptive knowledge developed by more effective individual learning, social learning results in larger brains. The Cumulative Cultural Brain Hypothesis predicts innovative ancestors—perhaps like the kind of individual innovativeness we see in chimpanzees (Reader et al., 2011).

Finally, environments have to be sufficiently rich (*λ*) to open the door to the regime of cumulative cultural evolution. Brains are costly, but this cost can be offset by more adaptive knowledge. The degree of mitigation is determined by *λ*. We find that higher *λ* values allow for the evolution of larger brains. Basically, you need to be in an environment where adaptive knowledge pays off well enough to pay for those costly brains.

One interesting, but speculative possibility that links these two parameters is that as the East African cradle of human evolution cooled and forests became savannah, our ancestors may have faced an increased selection for smaller brains helping to trigger the transition from asocial to social learning. That is, the forests were a richer ecology with higher *λ*, allowing for large-brained ancestors who could pay for their large brains through asocial smarts. As the forest thinned into savannah, the ecology became tougher and *λ* decreased, social learning may have provided a cheaper alternative to acquiring this knowledge and gaining more in order to maintain large brains in a calorie-poorer and less forgiving environment. Though we might infer such a scenario from our model, we would need to adjust these parameters within the model in order to test this hypothesis.

> Why some social learning is common but cumulative cultural evolution is rare

In addition to our main simulations that began with asocial learners, we also ran a set of simulations that began with social learners. Although social learning is widespread in the animal kingdom (Hoppitt & Laland, 2013) and the most realistic starting conditions are somewhere in- between these two extremes (no social learning and complete social learning), these realistic conditions are likely closer to no social learning than complete social learning. Nonetheless, running our simulations beginning with social learning provides an upper bound on our predicted patterns and also offers additional insights.

There are two key insights. The first is that social learning is maladaptive in a world with little knowledge (Figure 15). With little knowledge for social learners to exploit, asocial learners quickly invade. However, since some social learning is present, once sufficient knowledge has been generated, social learning is again at an advantage, with additional innovations generated in the process of social learning (Muthukrishna & Henrich, 2016). The second key insight is closely related: consistent with previous models (Boyd & Richerson, 1996), the presence of social learning expands the range of parameters in which cumulative culture is adaptive. Figure 9b shows a greater number of species with social learning (compared to Figure 9a). Figure 16a reveals that more monogamish societies are more likely to enter the realm of cumulative cultural evolution. Figure 16b reveals that cumulative cultural evolution is more likely to evolve when transmission fidelity is higher. Both Figures 16a and 16b reveal that the range of parameters that lead to the realm of cumulative cultural evolution expands if more social learning is present in the ancestral state.

**Figure 15.**
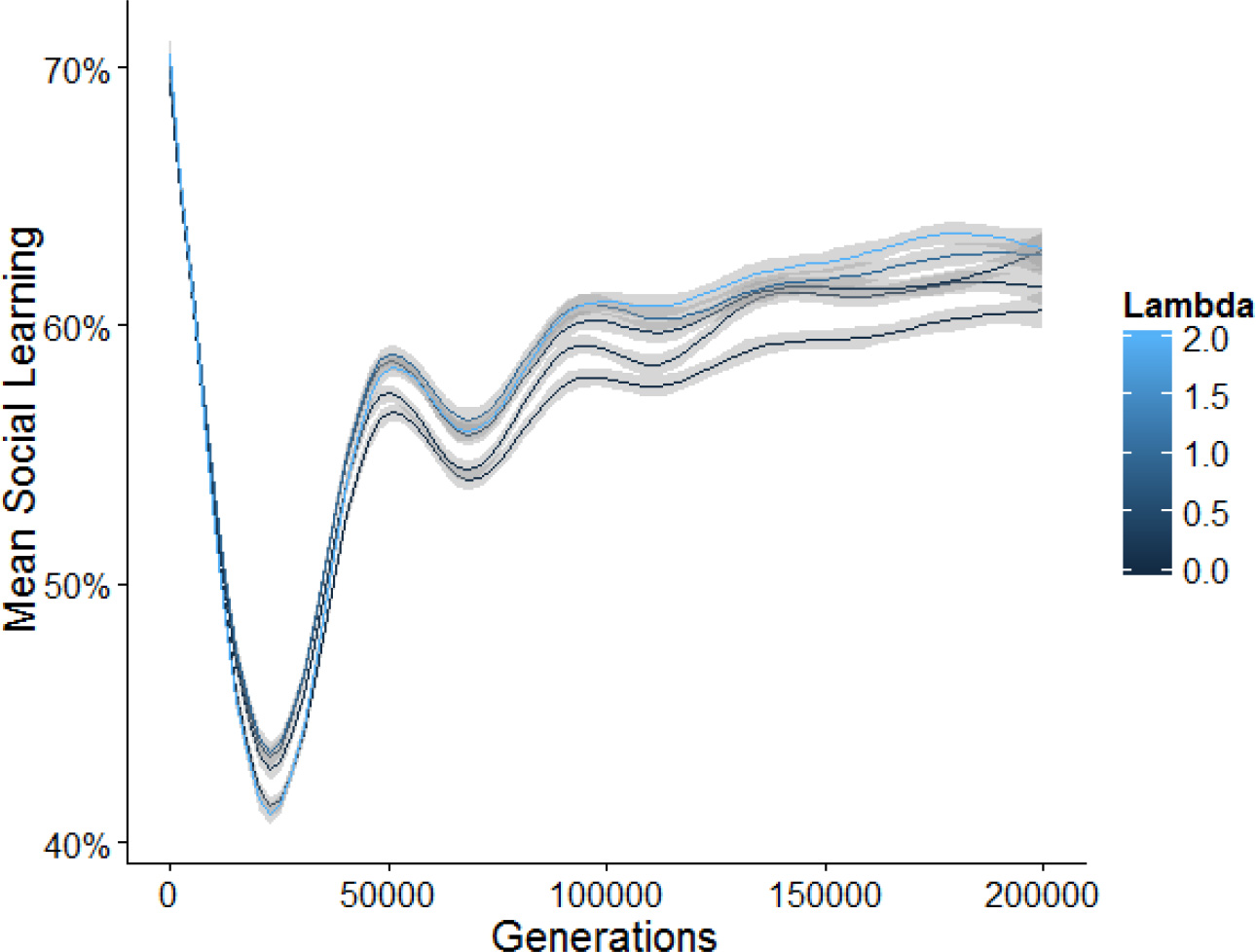
Social learning over generations starting with *s* = 1. 0. Social learning is maladaptive in the absence of adaptive knowledge. Asocial learners quickly invade. It is only when asocial learners have generated sufficient adaptive knowledge that social learners again have an advantage. Since we know that at least two regimes reliably emerge, mean social learning in these plots represents the relative number of conditions in which social and asocial learners emerge rather than a value of social learning characteristic of the world.

**Figure 16.**
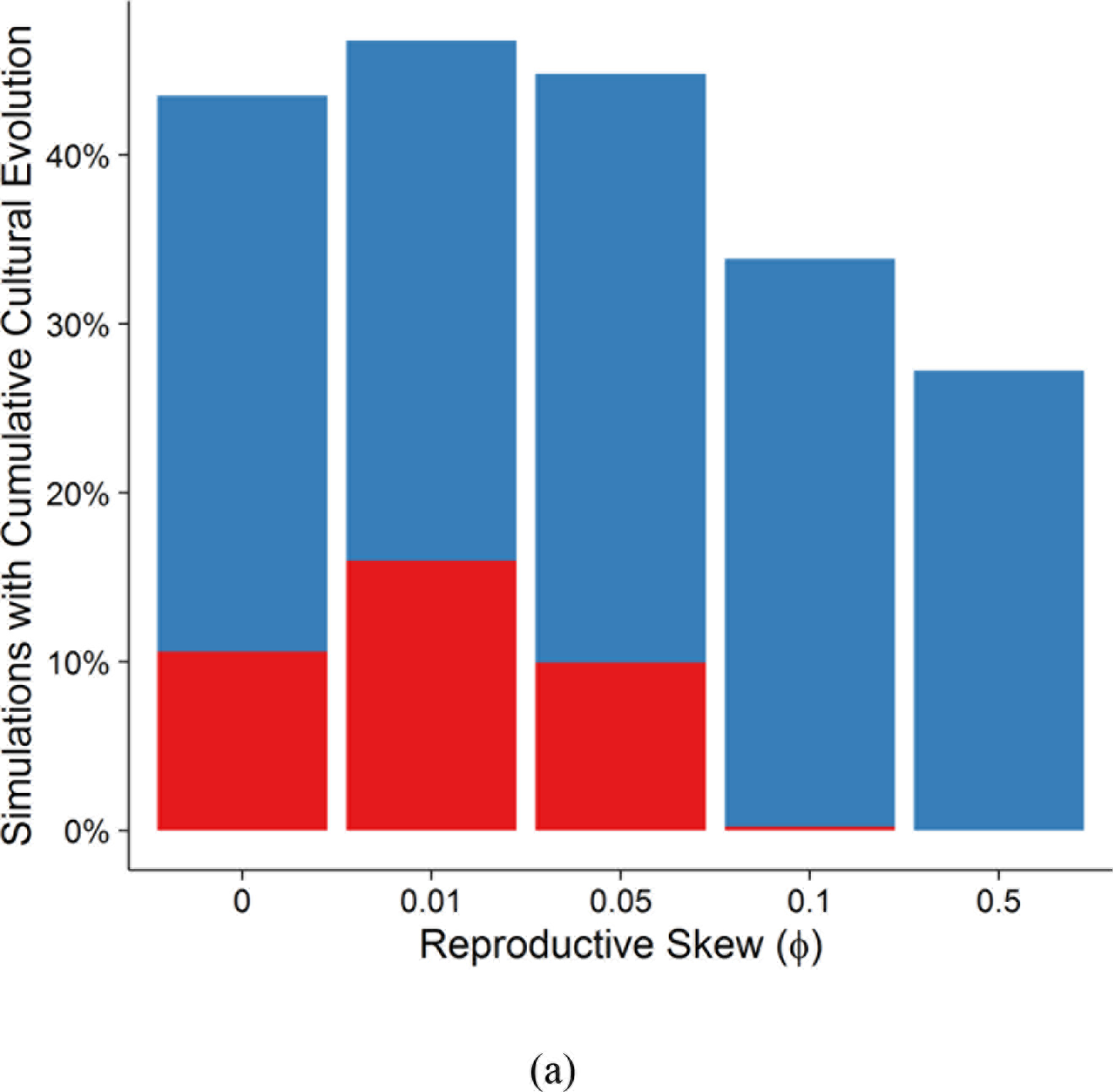
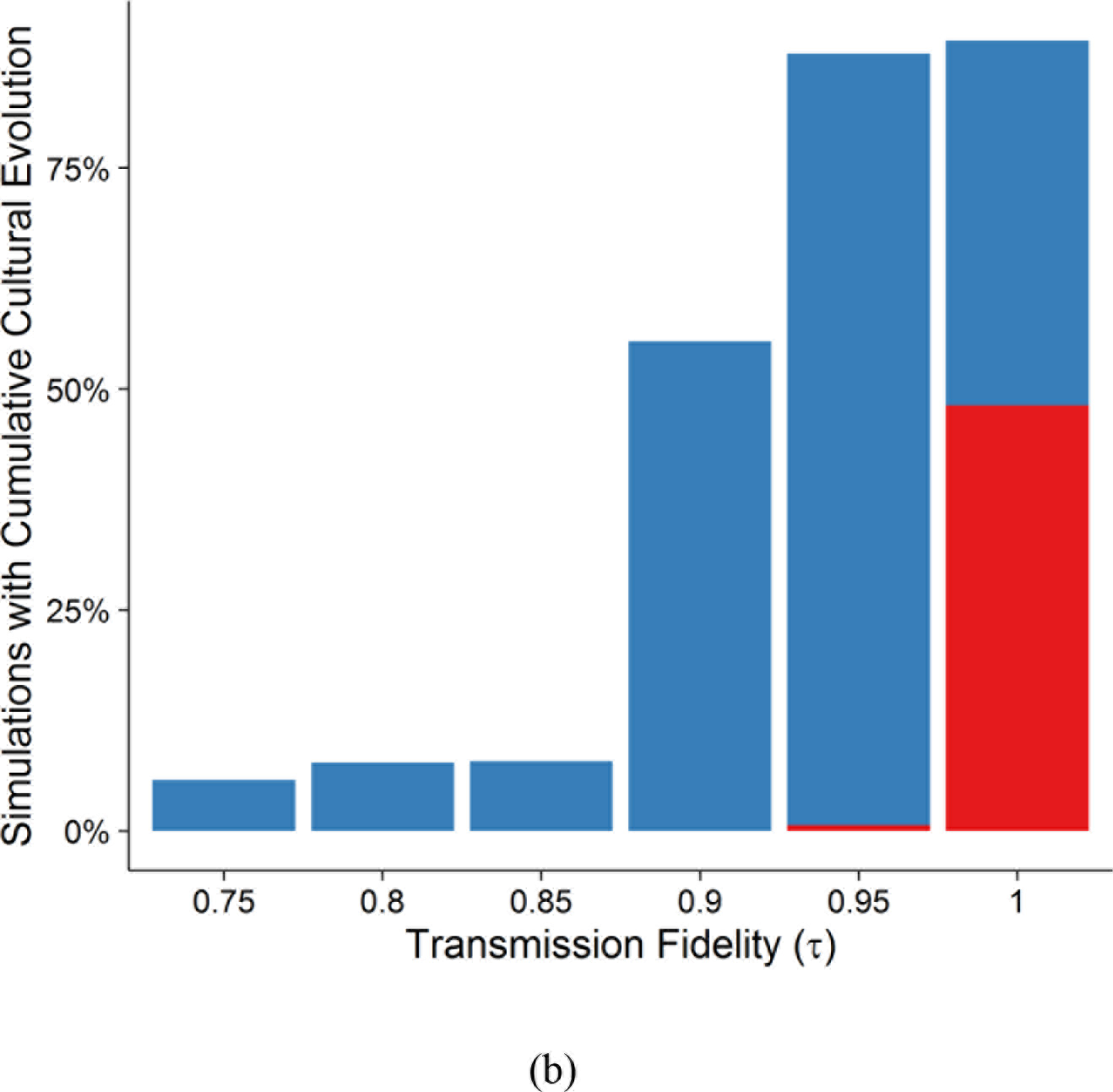
Percent of simulations in which cumulative cultural evolution evolves. Blue simulations are those that began with *s* = 1.0 and red simulations are those that began with *s* = 0.0. (a) across different values of reproductive skew (*ϕ*) and (b) across different values of transmission fidelity (*τ*).

## Discussion

In this discussion section we (1) summarize our key findings, (2) review these findings in the context of the cultural/general intelligence hypotheses and related work, and (3) discuss limitations of this work and ongoing inquiries.

### Summary of Key Findings

Our model provides a potential evolutionary mechanism that can explain a variety of empirical patterns involving relationships between brain size, group size, innovation, social learning, mating structures, and developmental trajectory, as well as brain evolution differences among species. It can also illuminate the different rates of evolution and overall brain size that have been found in different taxa and help explain why brain size correlates with group size in some taxa, but not others. In contrast to competing explanations, the key message of the Cultural Brain Hypothesis (CBH) is that brains are primarily for the acquisition, storage and management of adaptive knowledge and that this adaptive knowledge can be acquired via asocial or social learning. Social learners flourish in an environment filled with knowledge (such as those found in larger groups and those that descend from smarter ancestors), whereas asocial learners flourish in environments where knowledge is socially scarce, or expensive but obtainable through individual efforts. The correlations that have been found in the empirical literature between brain size, group size, social learning, the juvenile period, and adaptive knowledge arise as an indirect result of these processes.

The Cumulative Cultural Brain Hypothesis posits that these very same processes can, under very specific circumstances, lead to the realm of cumulative cultural evolution. These circumstances include when transmission fidelity is sufficiently high, reproductive skew is in a Goldilocks’ zone close to monogamy, effective asocial learning has already evolved, and the ecology offers sufficient rewards for adaptive knowledge. In making these predictions, the Cultural Brain Hypothesis and Cumulative Cultural Brain Hypothesis tie together several lines of empirical and theoretical research.

### Related Work

Under the broad rubric of the Social Brain or Social Intelligence Hypothesis, different researchers have highlighted different underlying evolutionary mechanisms (Dávid-Barrett & Dunbar, 2013; Gavrilets & Vose, 2006; McNally et al., 2012; McNally & Jackson, 2013). These models have had differing levels of success in accounting for empirical phenomena, but they highlight the need to be specific in identifying the driving processes that underlie brain evolution in general, and the human brain specifically. From the perspective of the CBH, these models have been limited in their success, because they only tell part of the story. Our results suggest that the CBH can account for all the empirical relationships emphasized by the Social Brain Hypotheses, plus other empirical patterns not tackled by the SBH. Moreover, our approach specifies a clear ‘take-off’ mechanism for human evolution that can account for our oversized crania, heavy reliance on social learning with sophisticated forms of oblique transmission (and possibly the emergence of adolescent as a human life history stage), and the empirically-established relationship between group size and toolkit size/complexity (Kline & Boyd, 2010)—as well as, of course, our species’ extreme reliance on cumulative culture for survival (Henrich, 2016).

Our simulation’s predictions are consistent with other theoretical work on cultural evolution and culture-gene coevolution. For example, several researchers have argued for the causal effect of sociality on both the complexity and quantity of adaptive knowledge (Kobayashi & Aoki, 2012; Powell, Shennan, & Thomas, 2009). Similarly, several researchers have argued for the importance of high fidelity transmission for the rise of cumulative cultural evolution (Enquist, Strimling, Eriksson, Laland, & Sjostrand, 2010; Henrich, 2004; Lewis & Laland, 2012).

Cultural variation is common among many animals (e.g., rats, pigeons, chimpanzees, and octopuses), but cumulative cultural evolution is rare (Boyd & Richerson, 1996; Henrich & Tennie, forthcoming). Boyd and Richerson (1996) have argued that although learning mechanisms, such as local enhancement (often classified as a type of social learning), can maintain cultural variation, observational learning is required for cumulative cultural evolution. Moreover, the fitness valley between culture and cumulative culture grows larger as social learning becomes rarer. Our model supports both arguments by showing that only high fidelity social learning gives rise to cumulative cultural evolution and that the parameter range to enter this realm expands if social learning is more common (see Figure 16). In our model, cumulative cultural evolution exerts a selection pressure for larger brains that, in turn, allows more culture to accumulate. Prior research has identified many mechanisms, such as teaching, imitation, and theory of mind, underlying high fidelity transmission and cumulative cultural evolution (Dean et al., 2012; Heyes, 2012; Morgan et al., 2015). Our model reveals that in general, social learning leads to more adaptive knowledge and larger brain sizes, but shows that asocial learning can also lead to increased brain size. Further, our model indicates that asocial learning may provide a foundation for the evolution of larger-brained social learners. These findings are consistent with Reader et al. (2011), who argue for a primate general intelligence that may be a precursor to cultural intelligence and also correlates with absolute brain volume.

The CHB is consistent with much existing work on comparative cognition across diverse taxa. For example, in a study of 36 species across many taxa, MacLean et al. (2014) show that brain size correlates with the ability to monitor food locations when the food was moved by experimenters and to avoid a transparent barrier to acquire snacks, using previously acquired knowledge. The authors also show that brain size predicts dietary breadth, which was also an independent predictor of performance on these tasks. Brain size did not predict group size across all these species (some of whom relied heavily on asocial learning). This alternative pathway of asocial learning is consistent with emerging evidence from other taxa. For example, in mammalian carnivores brain size predicts greater problem solving ability, but not necessarily social cognition (Benson-Amram, Dantzer, Stricker, Swanson, & Holekamp, 2016; Holekamp & Benson-Amram, 2017). These results are precisely what one would expect based on the Cultural Brain Hypothesis; brains have primarily evolved to acquire, store and manage adaptive knowledge that can be acquired socially or asocially (or via both). The Cultural Brain Hypothesis predicts a strong relationship between brain size and group size among social learning species, but a weaker or nonexistent relationship among species that rely heavily on asocial learning.

Our simulation results are also consistent with empirical data for relationships between brain size, sociality, culture, and life history among extant primates (e.g. Street, Navarrete, Reader, & Laland, 2017) and even cetaceans (Fox et al., 2017), but suggest a different pathway for humans. In our species, the need to socially acquire, store, and organize an ever expanding body of cultural know-how resulted in a runaway coevolution of brains, learning, sociality and life history. Of course, this hypothesis should be kept separate from the CBH: at the point of the human take-off, brain size may have already been pushed up by the coordination demands of large groups, Machiavellian competition, or asocial learning opportunities (Henrich, 2016). For example, Machiavellian competition may have elevated mentalizing abilities in our primate ancestors that were later high-jacked, or re-purposed, by selective pressure associated with the CCBH to improve social learning by raising transmission fidelity, thereby creating cumulative cultural evolution. Thus, the CBH and CCBH should be evaluated independently.

#### Synthesis and Naming

These ideas, which have been developed concurrently by researchers in different fields, are sufficiently new such that naming and labeling conventions have not yet converged. We use Cultural Brain Hypothesis and the Cumulative Cultural Brain Hypothesis for the ideas embodied in our formal model. We nevertheless emphasize that we are building directly on a wide variety of prior work that has used various naming conventions, including The Cultural Intelligence Hypothesis (Whiten & Van Schaik, 2007) and the Vygotskian Intelligence Hypothesis (Moll & Tomasello, 2007). And, of course, Humphrey (1976) originally described the importance of social learning in his paper on the social functions of intellect, though subsequent work has shifted the emphasis away from social learning and toward both Machiavellian strategizing and the management of social relationships. Whiten and Van Schaik (2007) first used the term “Cultural Intelligence Hypothesis” to argue that culture may have driven the evolution of brain size in nonhuman great apes. Later, Herrmann et al. (2007) used the same term to argue that humans have a suite of cognitive abilities that have allowed for the acquisition of culture. Supporting data for both uses of the term are consistent with the CBH and the CCBH (for a rich set of data and analyses, see Reader et al., 2011). We used two new terms not to neologize, but because though our approach is clearly related to these other efforts, our approach contains novel elements and distinctions not clarified or formalized in earlier formulations.

### Simplifications, Extensions and Future work

Note that our model seeks to (1) show why brain size, adaptive knowledge, social learning, group size, and lifespan are intercorrelated across the animal kingdom (CBH) and (2) how the very same processes that lead to these interconnections, can, under some specific circumstances, lead to the realm of cumulative cultural evolution—the uniquely human pathway. Within the realm of cumulative culture, the dynamics change in ways that are not captured by this model. For example, in order to sustain ever-growing levels of cultural complexity, cultures can generate ways to increase sociality and transmission fidelity. With sufficiently complex culture, mechanisms may evolve to more efficiently share the fruits of rare innovations, allowing for increases in cultural variance that may be individually costly. Moreover, cumulative culture, once acquired, can increase an individual efficacy in subsequent asocial learning (for a discussion of these ideas, see Muthukrishna & Henrich, 2016).

In developing the simulation, we formalized the minimal set of assumptions and parameters that capture the logic of the CBH and CCBH. There are a number of extensions, variations, and additional parameters that would improve our understanding of the evolution of brain size.

There were several assumptions that simplified our model, making it more computationally tractable. Future models may address some of these shortcomings and explore additional parameters. One such improvement is to explicitly track different cultural traits with different cognitive costs and fitness payoffs. By doing this, we could better explore the benefits to migration and cultural recombination. We would also like to more fully explore the impact of the relationship between adaptive knowledge and carrying capacity. Currently, the richness of the ecology only affects individual survival based on paying the calorie cost of costly brains, but the richness of the ecology also affects the carrying capacity of the population with consequent effects for the dynamics between brain size, adaptive knowledge and population size.

Another previously mentioned future improvement is the endogenization of transmission fidelity (*τ*) and reproductive skew (*φ*). These parameters are themselves subject to genetic and cultural evolutionary processes and thus ought to be modeled as endogenous variables. In our model, we can discuss the effect of different evolutionary outcomes or values of transmission fidelity and reproductive skew, but not their evolution.

Two or three regimes emerged in our models based on different ecological and phylogenetic constraints. In a future model, we plan to explore the adaptive dynamics of these different regimes, exploring the invasion fitness of the different equilibrium states discovered in our model. These models will help us better understand the evolutionary dynamics that may have occurred when different previously geographically separated hominin species encountered each other (e.g., the European encounter between modern humans and their larger-brained Neanderthal cousins).

The key improvements that we are eager to explore could be summarized as: (1) endogenizing the evolution of transmission fidelity and reproductive skew, (2) explicitly tracking different cultural traits with different cognitive costs and fitness payoffs, and (3) more thoroughly exploring the brain shrinkage that occurs during the transition from reliance on asocial learning to reliance on social learning. These results hint that the process underlying the Cultural Brain Hypothesis and Cumulative Cultural Brain Hypothesis may also help explain evidence suggesting that human brains have been shrinking in the last 10,000 to 20,000 years (Ruff, Trinkaus, & Holliday, 1997). Although this shrinkage in brain size corresponds to shrinking in body size, it may be evidence that our species is not at equilibrium.

## Acknowledgements

The computational model was enabled in part by support provided by Westgrid and Compute Canada. J.H. acknowledges support from the Canadian Institute for Advanced Research.

## Footnotes

i Curves generated using Magnusson (2016) (rpsychologist.com)

